# Social Functioning in Autism: A Systematic Review and Meta-analysis

**DOI:** 10.64898/2026.03.20.713084

**Authors:** Siyi Li, Huanqing Wang, Guoqiu Chen, Xi Cheng, Yuxi Wang, Wanyan Hu, Antonia Hamilton, Leonhard Schilbach, Li Yi, Kaat Alaerts, Zhanjun Zhang, Xi-Nian Zuo, Yin Wang, Yinyin Zang

## Abstract

Atypical social functioning is a core feature of autism, yet findings remain fragmented across components and development. We aimed to systematically integrate this literature and characterize the organization, development, and moderators of social functioning in autism. We conducted a systematic review and meta-analysis of behavioral studies published between January 1990 and August 2025, identified through PubMed, Web of Science, and prior reviews, including studies with clinically diagnosed autistic individuals and neurotypical controls. A qualitative synthesis and two complementary quantitative meta-analyses were performed, with risk of bias evaluated through study-level characteristics. A total of 2,622 studies (94,114 autistic and 172,847 neurotypical individuals across 32 countries) were included, covering 22 social components that clustered into five domains. Overall group differences were substantial (Hedges’ *g* = -0.744, 95% CI [-0.797, -0.690]). Differences emerged earliest in motivation-based processes (∼6 months), followed by motor, emotion, and inference domains, and showed age-related divergence alongside improvement in some skills. Cross-domain analyses revealed stronger interdependencies in autism and an organizational pattern most consistent with serial relationships among domains. These findings should be interpreted in light of methodological heterogeneity, underpowered samples, and uneven cultural representation. Together, the results provide an integrative framework for understanding the organization and development of social functioning in autism, with implications for precision subtyping, developmentally timed interventions, and neurodiversity-informed research and policy. This study was pre-registered (PROSPERO: CRD42024566141).

## Introduction

Atypical social functioning is a core feature of autism spectrum condition (ASC), affecting around 1% of the global population and impacting communication, relationships, and social navigation across the lifespan ^1–3^. Despite decades of research, interventions for social difficulties in autism have shown limited real-world efficacy. While some therapies improve specific social skills in structured settings, these gains often fail to generalize to everyday interactions, and their long-term sustainability remains unclear ^4,5^.

This lack of progress reflects a persistent gap between basic research and clinical practice—a challenge amplified by the complexity and heterogeneity of autism ^6,7^. Basic research often isolates individual aspects of social functioning, adopting a reductionist approach that overlooks the broader, interdependent nature of social abilities in autism ^8,9^. Clinically, interventions focus primarily on observable social performance—such as verbal communication and nonverbal behaviors—while underlying cognitive mechanisms that shape these abilities remain underexplored ^10^. However, social difficulties in autism are neither static nor isolated but rather part of a dynamic system of interrelated abilities, spanning basic processes (e.g., social attention, imitation) to higher-order skills (e.g., theory of mind, social communication) ^11,12^. Here, we use the term social functioning to encompass this full range—from foundational socio-cognitive and motivational processes to observable social behaviors—as all contribute to how individuals engage with the social world ^92^. Despite growing recognition of these complexities, research remains fragmented and poorly integrated, making it difficult to translate scientific advances into clinical applications. As a result, critical questions—which aspects of social functioning show the greatest divergence from neurotypical (NT) expectations and what mechanisms drive these differences—remain unresolved, limiting the development of targeted supports tailored to different social profiles.

Critical methodological challenges continue to limit progress in understanding social functioning in autism. Issues such as underpowered samples, unbalanced age distributions, and skewed sex ratios have led to inconsistent findings ^13–15^. Additionally, the widespread use of artificial experimental tasks often fails to reflect the complexity and variability of real-world social interactions, reducing the clinical and practical relevance of laboratory-based research ^16–18^. Compounding these challenges, different studies focus on different aspects of social functioning, making it difficult to integrate findings into a cohesive framework ^19,20^. Even within the same domain, studies use varying methodologies, and even when employing the same methodology, they often rely on different measurement indices (e.g., multiple eye-tracking metrics have been adopted for social attention studies ^21^). These inconsistencies contribute to conflicting conclusions and constrain efforts to identify meaningful patterns that could improve autism interventions and better align scientific inquiry with the lived experiences of autistic individuals.

Developmental trajectories add further complexity to the understanding of social functioning in autism. Different aspects of social abilities emerge at different ages and can change across the lifespan^1^. For instance, foundational skills such as imitation may generally improve with age, while more complex abilities, like perspective-taking and social reciprocity, often become increasingly challenging during adolescence and adulthood as social interactions become more nuanced and demanding ^22–24^. These age-specific patterns suggest that there may be critical windows where targeted support could be most effective. However, many existing approaches do not fully account for developmental change, often applying the same interventions across all ages^6^. Understanding how social abilities evolve over time is essential for optimizing intervention timing and improving long-term outcomes ^25,26^.

Theoretical and empirical models suggest that social cognition is organized hierarchically, with lower-level processes scaffolding the development of more complex social abilities. For instance, Malle (2020) ^27^ proposed a tree-like hierarchical model in which foundational skills—such as social attention, imitation, and action understanding—support the development of higher-order capacities like mentalizing, social reasoning, and moral judgment. Supporting this view, Schurz et al. (2021) ^28^ conducted a large-scale meta-analysis of neuroimaging studies and identified a functional gradient in social cognition, showing that brain regions involved in low-level mirroring processes are systematically distinct yet interconnected with those supporting high-level inferential functions. Beyond these structural models, developmental theories such as the social motivation hypothesis ^29^ suggest that early atypicalities in social orienting and reward sensitivity may contribute to later challenges in joint attention, empathy, and communication. This cascading view aligns with broader theoretical frameworks highlighting the developmental interdependence of social cognitive processes ^12^. Longitudinal findings further support this idea, indicating that early social attention and motivation can predict later social abilities, including language and theory of mind ^30–32^. Taken together, these frameworks suggest that social abilities may become apparent at different developmental periods and are interdependent and developmentally scaffolded—an insight that motivates our effort to map their organization in autism more comprehensively. Here, we examine cross-sectional patterns across ages to explore potential ordering and interdependence among social abilities, while acknowledging that such patterns cannot establish causal developmental sequences. Our goal is not to test developmental order per se, but rather to evaluate whether observed inter-domain dependencies in autism reflect organizational patterns consistent with these well-supported theoretical frameworks.

Although autism is one of the most heritable neurodevelopmental conditions^33^, social behavior is also shaped by sociocultural norms, which influence both the expression of autism-related traits and how these behaviors are perceived ^34,35^. What is considered socially appropriate, atypical, or unexpected varies widely across cultures, affecting diagnostic practices, intervention approaches, and lived experiences. However, research on social functioning in autism has largely prioritized biological mechanisms, often overlooking the critical role of cultural context^36^. While there is growing recognition that cultural norms shape social behaviors and perceptions of autism, empirical data on these influences remain limited^37–39^. Systematic research is needed to determine how sociocultural factors interact with autistic social development, identify global patterns and cultural variations, and develop more inclusive, contextually relevant models of social functioning^40,41^. Additionally, recent perspectives on neurodiversity emphasize valuing differences rather than enforcing conformity to social norms, reinforcing the need for culturally responsive frameworks that support diverse styles of social interaction.

Despite these challenges, no integrated synthesis has integrated findings across all major aspects of social functioning in autism while accounting for methodological variability, developmental change, and sociocultural influences^6^. Prior meta-analyses have typically focused on isolated components, limited age ranges, or single-context designs, leaving the field without an integrative framework that reflects the full complexity of autistic social experience. To address this gap, we adopted a multi-layered analytic strategy. First, we conducted a qualitative systematic review to map the breadth and distribution of reported social differences across 22 social components. Second, we performed a quantitative meta-analysis of group differences to estimate the magnitude of autistic–neurotypical differences and to examine developmental, methodological, and sociocultural moderators. Third, recognizing that social abilities are interdependent rather than isolated, we conducted a quantitative meta-analysis of behavioral correlations, including meta-analytic structural equation modeling, to evaluate how different domains of social functioning relate to one another and whether these relationships conform to theoretically proposed organizational architectures. Together, these complementary analyses provide a structured framework for understanding how social abilities are organized, develop over time, and are shaped by contextual factors in autism, with implications for future research, developmentally timed interventions, and precision-based clinical applications ^42,43^.

## Result

### Study Characteristics and Scope

Based on prior theoretical frameworks (see details in *Methods* section) ^44–47^, a systematic review identified 2,622 eligible studies published between January 1990 and August 2025 from PUBMED, Web of Science, and previous meta-analyses. The review involved data from 94,114 autistic and 172,847 neurotypical individuals across 32 countries. The studies examined 22 key social components ^48^ (e.g., social attention, action perception, person perception, empathy, theory of mind, self-processing, imitation, pretend play, deception, communication, see the complete list in **Fig. 1a**). Definitions of these components are provided in **Extended Data Table 1**. The PRISMA flowchart **(Fig. 2)** illustrates the study selection process. As the present review focused on social difficulties without intellectual disability, studies on autistic individuals with co-occurring intellectual disabilities were excluded. This meta-analysis was pre-registered on PROSPERO (see details in Methods).

**Fig 1.**
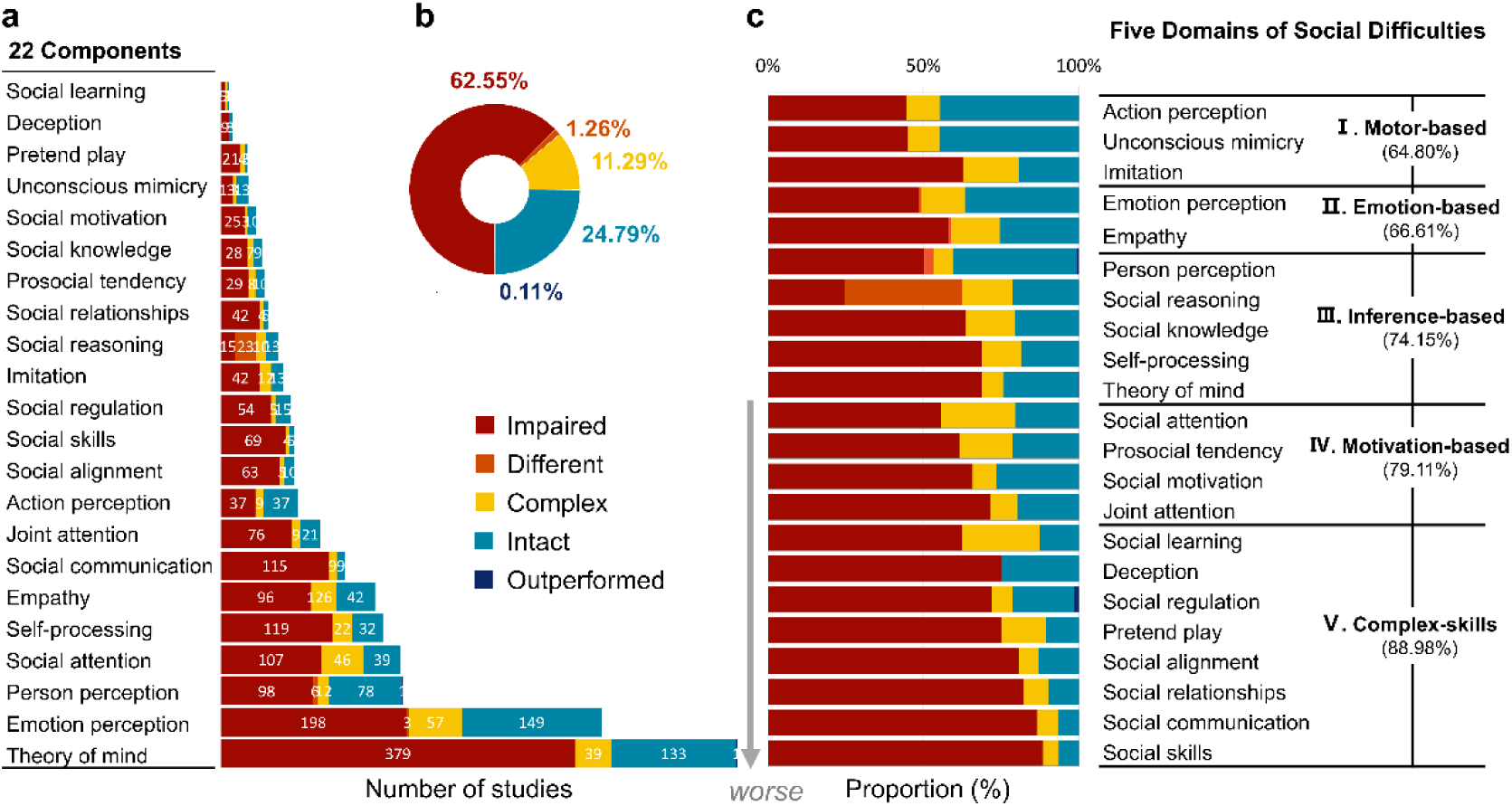
Overall social functioning in autism. **(a)** Study distribution across 22 social components. Each study was coded into one of five categories based on its main conclusion: Impaired (i.e., poorer performance in ASC), Different (i.e., different performance without superiority/inferiority), Complex (i.e., varying group differences across conditions), Intact (i.e., comparable performance with NT populations), or Outperformed (i.e., better performance in ASC). **(b)** Proportion of conditions across all studies. Overall, 75.21% of the studies reported alterations (i.e., impaired + different + complex + outperformed) in the autistic group. **(c)** Prevalence of alterations in each component, ranked as the proportion of each category. Guided by prominent neurocognitive theories in autism (see details in Methods section), we grouped social difficulties into five domains (I-V), with motor-based components as impacted the least and complex skills as impacted the most. The percentage figures in parentheses represent the reported alterations averaged across components in each domain. Panel (c) summarizes a qualitative classification of study outcomes. This five-category system was not used in any of the quantitative meta-analyses (e.g., effect size calculations, moderator analyses, or correlation tests), which were conducted solely using continuous effect size metrics (see ‘Selection criteria and analytical strategy’ in *Methods*).

**Fig. 2.**
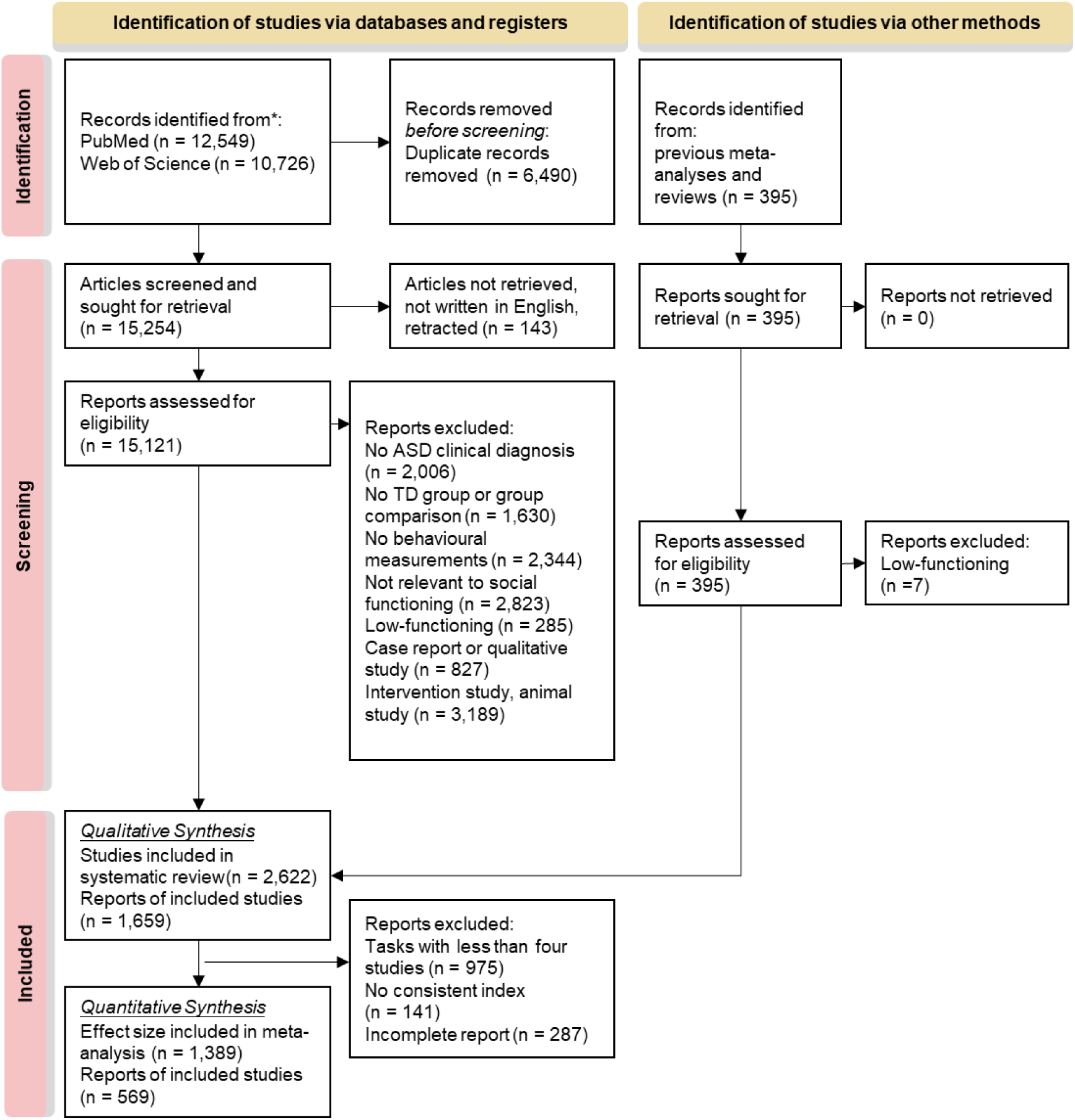
PRISMA flow chart of the systematic review process.

### Methodological Scrutiny

Significant methodological variability and biased distributions were observed across studies. Research efforts were unevenly distributed across social components **(Fig. 1a)**, with theory of mind having been extensively studied (552 studies), whereas social learning was notably underrepresented (8 studies). The ages of autistic participants ranged from 0.5 to 54.6 years (median = 13.41 years), displaying a bimodal distribution **(Extended Data Fig. 2b)**. Sex ratios were highly skewed, with a median male ratio of 0.833 **(Extended Data Fig. 2c)**. Sample sizes varied widely, ranging from 3 to 1,905 participants (median = 22) **(Extended Data Fig. 2a)**. Power analyses indicated that 69.957% (971/1388) of studies did not meet the minimally required sample size **(Extended Data Table 3)**, increasing the risk of false-positives and reduced reproducibility, though underpowered studies did not significantly inflate effect sizes (*p* = 0.926) **(Extended Data Fig. 4)**.

Four major sources of methodological heterogeneity influenced the reported social differences in autism **(Fig. 3a)**. First, self-report surveys yielded larger social differences than experimental tasks (t = 7.477, *p* < .001), raising concerns about subjective biases. This is consistent with prior findings of weak convergence between survey and experiment ^49^. Second, differences in the experimental paradigms used significantly impacted findings. For example, two task paradigms have been used for studying social motivation: social seeking—the tendency to preferentially select social over non-social stimuli in simultaneous-choice paradigms—and social reward, defined as behavioral enhancement in response to socially rewarding outcomes (e.g., increased accuracy or speed when a smiling face is used as a reward). We found significant group differences in social seeking (Hedges’ *g* = -0.735, 95% CI [-1.453, -0.016]) but not in social reward (Hedges’ *g* = 0.025, 95% CI [-0.054, 0.103]; t = 7.439, *p* < .001), indicating that while autistic individuals show reduced preference for social stimuli, they do not differ in reward-driven task performance when the reward is social. This dissociation is consistent with prior meta-analyses (see direct comparison in **Supplementary Fig. 2**) and underscores the value of task-specific characterizations of social functioning^50^. Third, the choice of performance indices also influenced results, as reaction time measures revealed greater differences than accuracy measures in action perception tasks (t = 2.373, *p* = 0.024). Finally, ecological validity played a crucial role, with naturalistic free play detecting more pronounced social differences than structured laboratory tasks (rho = -0.460, *p* = 0.012). Notably, task paradigms significantly influenced reported social differences in empathy, self-processing, person perception, and social motivation but had minimal influence on theory of mind (ToM), emotion perception, joint attention, and imitation **(Fig. 3b)**. We also examined whether effect sizes varied by reporter (self vs. proxy) and no significant moderation was observed (t (209) = 1.194, 95% CI = [-1.447, -1.076], *p* = 0.234). Although previous studies have reported discrepancies between self- and proxy-reports in autism ^51,52^, we did not detect such an effect in our meta-analysis. Given that fewer than 10% of studies used proxy reports and that reporter type was strongly confounded with age group, this null result should be interpreted cautiously. It is possible that true discrepancies exist but are masked at the aggregated level due to heterogeneity across tasks, populations, and contexts.

**Fig. 3.**
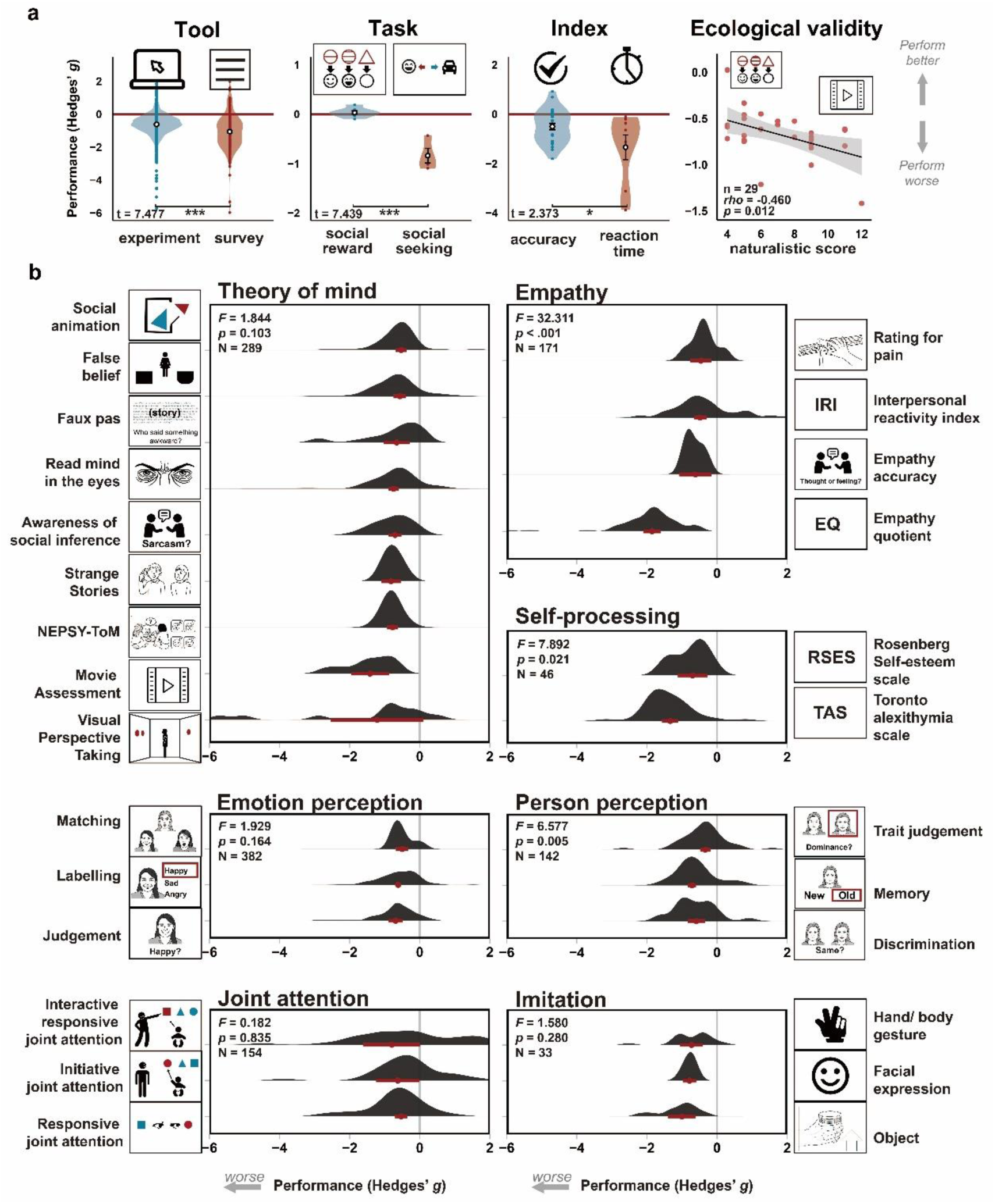
Methodological heterogeneity and its effects on reported social differences in autism. **(a)** Four major sources of methodological variabilities in autism research that had significant effects on the degree of social difficulties: tools (e.g., more alterations observed in surveys than experiments, independent samples t(1386) = 7.477, 95% CI = [0.355, 0.607], η^2^ = 0.039, *p* <.001), task paradigms (e.g., atypical social motivation was only observed in social seeking paradigm, but not social reward paradigm, t(9) = 7.439, 95% CI = [0.606, 1.135], η² = 0.860, *p* <.001), indices (e.g., plot-light task performance of action perception was worse when analyzing reaction time variables than accuracy variables, t(33) = 2.373, 95% CI = [0.119, 1.556], η² = 0.146, *p* = 0.024), and ecological validity (e.g., more social differences detected by naturalistic/interactive games than tightly controlled lab tasks, *p* = 0.012). Hedges’ *g* values on the y-axis represent altered performance (i.e., ASC-NT difference, more negative values indicate poorer performance in ASC). While dots indicate the estimated effect size; Red and blue violin plots represent greater and lesser alteration, respectively; the bounds of the box demonstrates the ±1 standard error(SE); \**p* < 0.05; \*\**p* < 0.01; \*\*\**p* < 0.001; The shaded area in the line plot represents the 95% confidence interval. **(b)** Alteration degree and density distribution for each task paradigm. Only components with multiple task paradigms (and meet the criteria for effect size calculation) were shown here. Red dots and error bars indicate the estimated effect sizes and 95% CI, respectively. All statistical tests were two sided.

Publication bias was detected in several tasks based on funnel plot asymmetry **(Extended Data Fig. 3)**. Rightward asymmetry in tasks such as the Toronto Alexithymia Scale (TAS), false belief task, and emotion labeling task suggested overestimated effect sizes. Some tasks, such as visual perspective-taking and emotion judgment, showed non-significant Egger’s test results, suggesting either high heterogeneity or a genuine absence of bias **(Supplementary Table 4)**.

### Group-difference meta-analysis

Across all 22 social components, 75.21% of studies reported differences in social functioning in autism, while 24.79% found no significant group differences **(Fig. 1)**. For a qualitative systematic review, study conclusions were categorized into five groups: “Impaired” (62.55%, indicating poorer performance in autistic individuals), “Different” (1.26%, distinct but not necessarily better or worse), “Complex” (11.29%, varying group differences depending on conditions), “Intact” (24.79%, comparable performance to neurotypical populations), and “Outperformed” (0.11%, superior performance in autistic individuals). A quantitative meta-analysis on effect size confirmed substantial group differences (Hedges’ *g* = -0.744 [-0.797, -0.690], *p* < .001), with findings remaining stable after excluding small-sample studies (**Supplementary Fig. 1**). These results were highly consistent with previous meta-analyses on individual social components (**Supplementary Fig. 2**), reinforcing the validity of our findings against the established literature.

Using a theory-driven framework (see details in *Methods* section), social differences in autism were organized into five domains, reflecting a progression from mild to more pronounced differences (**Fig. 1c**). Motor-based differences (e.g., action perception, imitation) were the least pronounced, followed by emotion-based (e.g., emotion recognition, empathy) and inference-based domains (e.g., ToM, social reasoning). Motivation-based differences (e.g., social attention, social motivation) were reported as more severe, while complex social skills (e.g., social communication, relationship-building) exhibited the largest overall differences.

To estimate the earliest observed social differences, we analyzed the reported age of onset for each domain (**Fig. 4a**): Motivation-based differences (e.g., joint attention) were the earliest to appear, with reports as early as 6 months of age^53^, followed by motor-based differences (i.e., imitation) at 12 months^54^, emotion-based differences at 18.2 months (i.e., empathic concern) ^55^, and inference-based differences at 36.8 months (i.e., implicit theory of mind) ^56^. This sequence of ‘motivation → motor → emotion → inference’ closely follows the typical developmental pattern of social functioning in neurotypical children^27^, where foundational social abilities emerge first and provide the basis for more complex social skills^57^.

**Fig. 4.**
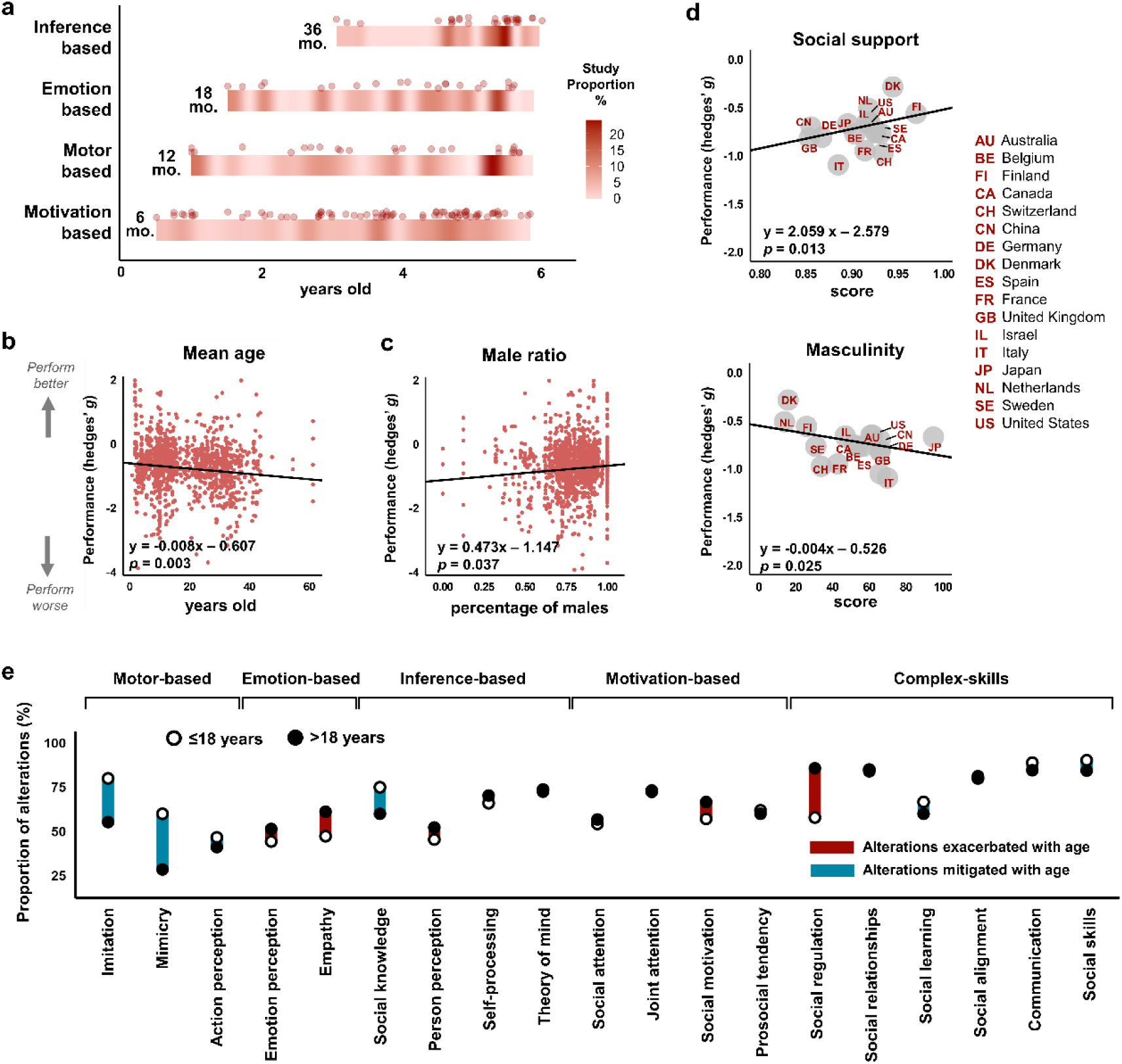
Onset, dynamics, and moderators of social difficulties in autism. **(a)** Developmental onset of social difference reported in each domain. Red dots represented each study and the density heatmap below showed the distribution of studies showing an effect across early childhood development (infancy and preschool age). The earliest observed group differences in autism, based on available cross-sectional data, are in motivation (∼6 months), followed by motor (∼12 months), emotion (∼18.2 months), and inference (∼36.8 months). These age estimates should be interpreted as the earliest ages at which group differences have been reported in literature, not as evidence of a within-individual longitudinal developmental trajectory. Meta-regression analyses are performed for **(b-c)** demographic factors (i.e. mean age and male ratio) of autistic samples, and **(d)** sociocultural moderators (i.e., perceived social support and masculinity in the country of residence). Hedges’ *g* values on the y-axis represent altered performance (i.e., ASC-NT difference, more negative values indicate poorer performance in ASC). Regression lines and p-values indicate the significance of correlations after hyperbolic and symmetric Gaussian transformation. **(e)** Changeability and dynamics of social difficulties across time. Proportion of reported alterations (y-axis) by component (x-axis) across five domains, comparing period of childhood (≤18 years, open circles) versus adulthood (>18 years, filled circles). Red lines indicate social difficulties become significantly worsening in adulthood, whereas blue lines indicate mitigation in adulthood.

### Cross-domain correlation meta-analysis

To examine how different domains of social functioning relate to one another, we conducted the correlation analyses in two stages. First, we performed an exploratory analysis comparing the strength of cross-domain correlations between autistic (ASC) and neurotypical (NT) groups, asking whether associations among domains differed in magnitude between groups. Second, we used meta-analytic structural equation modeling (MASEM) to evaluate whether these correlations were organized according to alternative theoretically motivated architectures. This two-stage approach allows group differences in association strength to be distinguished from differences in higher-order organizational structure.

In the exploratory analysis, we found that autistic individuals exhibited stronger cross-domain correlations than neurotypical individuals (*F* (1, 98.2) = 8.278, partial R^2^ = 0.078, 95% CI = [0.008, 0.201], *p* = 0.005), especially between emotion-based and inference-based domains (*F* (1,25.806) = 10.385, *p* = 0.034), suggesting tighter coupling among social (**Extended Data Fig. 1a & 1b**).

To further examine the organizational structure of these interdependencies, we tested whether the observed correlations were most consistent with a cascading architecture proposed in prior literature ^12^. Building on established developmental theories ^27^ , as well as our observation that different social differences are first reported at different ages in aggregated cross-sectional data (**Fig. 4a**), we evaluated whether more foundational domains were structurally associated with more complex domains in a manner consistent with cascading models of social functioning. We applied meta-analytic structural equation modeling (MASEM) to compare three theory-driven directional architectures ^12,27^: a *Serial* model, in which influence proceeds sequentially from motivation- to motor-, emotion-, inference-based domains and ultimately to complex social skills; a *Radial* model, in which motivation-based processes directly influence all other domains; and a *Network* model allowing all pairwise influences. Model comparison favored the Serial model, which showed the best fit (lowest BIC) (**Extended Data Fig. 1c**; **Supplementary Data Table 6**). A permutation test validated its superiority over 120 alternative unidirectional permutations (3/120, *p* = 0.025), and a likelihood ratio test revealed significant group differences in model paths (*p* = 0.023). Notably, autistic individuals showed stronger paths from emotion-based to inference-based domains and from inference-based domains to complex social skills (**Extended Data Fig. 1c**, red arrows). While the MASEM results support the Serial model as best fitting the observed cross-domain correlations, this finding does not imply a fixed developmental sequence or causal influence, but rather reflects an organizational pattern consistent with prior developmental theory.

### Moderating Factors Influencing Social Functioning

Moderating effects analysis identified several key factors influencing social functioning in autism. Overall, social performance declined with age (*β* = -0.008, *p* = 0.003, **Fig. 4b**). Due to data scarcity and heterogeneity, we were unable to delineate reliable developmental patterns for each individual social component. Instead, given the bimodal age distribution in the literature (**Extended Data Fig. 2b**), we compared childhood and adulthood to examine the developmental shifts and dynamics of social functioning in autism (**Fig. 4e**). Two distinct patterns emerged: emotion-based, inference-based, and motivation-based components, as well as certain complex social skills (e.g., social regulation and social relationships), declines with age. In contrast, motor-based components and some complex social abilities (e.g., social alignment, social learning, and social communication) improve in adulthood. Sex ratios in study samples also appeared to influence reported outcomes. A higher proportion of males in autistic samples was associated with better reported social functioning (*β* = 0.473, *p* = 0.037, **Fig. 4c**), suggesting that sex-related factors may contribute to variability in social outcomes.

Sociocultural factors also played a role (**Supplementary Table 5**). Higher perceived social support in a country was associated with smaller differences in social functioning between autistic and neurotypical individuals (*β* = 2.020, *p* = 0.003), while higher national masculinity (i.e., societal emphasis on achievement and competition) was linked to greater differences in social functioning (*β* = -0.003, *p* = 0.047) (**Fig. 4d**). Other factors, including participant characteristics (IQ scores, diagnostic criteria, comorbidities), study attributes (sample size, publication year, journal impact factor, citations), and national indicators (incidence of autism, economy, democracy, etc.), showed no significant effects (**Extended Data Fig. 4** and **Supplementary Table 5**).

## Discussion

This study presents the largest and most systematic synthesis of social functioning in autism, integrating findings from over 2,500 empirical studies spanning 35 years. Unlike previous meta-analyses, which focus on single social components in isolation, our study uncovers the interconnected nature of social difficulties in autism, showing how 22 social components cluster into five hierarchical domains—motor-based, emotion-based, inference-based, motivation-based, and complex skills **(Fig. 1c)**. Crucially, this study provides new insights into: (1) the hierarchical organization of social functioning, consistent with theoretical models suggesting that lower-level processes scaffold the development of higher-order abilities^12,27,28^; (2) age-related patterns in group differences and their malleability across development **(Fig. 4)**; (3) stronger cross-domain behavioral correlations in autism **(Extended Data Fig. 1)**, suggesting tighter coupling among social processes; and (4) methodological and sociocultural influences that may account for prior inconsistent findings **(Fig. 3 & 4)**. These findings advance understanding of autism’s complex social profiles and provide a foundation for developmentally tailored and culturally informed public health strategies. Importantly, this work also advances our understanding of the underlying structure and development of human social cognition more broadly, revealing how different social capacities emerge, interact, and diverge across individuals and populations.

Our results highlight a potential ordering of social functioning differences across age groups, wherein differences in social motivation are observed earlier in life than differences in more complex abilities (**Fig. 5**). While our meta-analysis does not rely on longitudinal datasets and cannot establish causality, the integration of multiple lines of evidence—including the early reported motivation-based differences (**Fig. 4a**), their larger proportion of difficulties (**Fig. 1c**), and the downstream pathways identified by a new MASEM analysis (**Extended Data Fig. 1c**) —suggests that social motivation may play a pivotal role in shaping later developmental outcomes. This interpretation aligns with both developmental theories and empirical findings. Theoretical models propose that foundational social processes, such as motivation and joint attention, enable infants to engage with caregivers and learn from social input^12,29^. Empirical evidence supports this view: for example, infants’ gaze following at 10 months predicts mental-state language use at 2.5 years and ToM at 4.5 years^31^; social motivation between 6 and 12 months correlates with joint attention skills at 15 months and language abilities at 24 months, which is important for developing social communication skills^32^; and longitudinal studies show that infants later diagnosed with ASC exhibit declined social motivation as early as 2 months^30,58^. Moreover, a series of intervention studies also use declined social engagement as an early detection index and perform effective interventions to improve high-risk infants’ social skills^58,59^. Taken together, these findings are consistent with a functional cascade^12,29^, whereby initial differences in motivation may limit opportunities for interactive learning, subsequently influencing downstream domains such as emotion understanding, inference-making, and complex social skills. By mapping interdependencies among social processes (**Fig. 5** and **Extended Data Fig. 1**), our results offer a scaffold for evaluating and timing interventions more precisely—especially in early childhood, when motivational and motor-based skills are still forming and may influence the developmental pathways of downstream abilities.

**Fig. 5.**
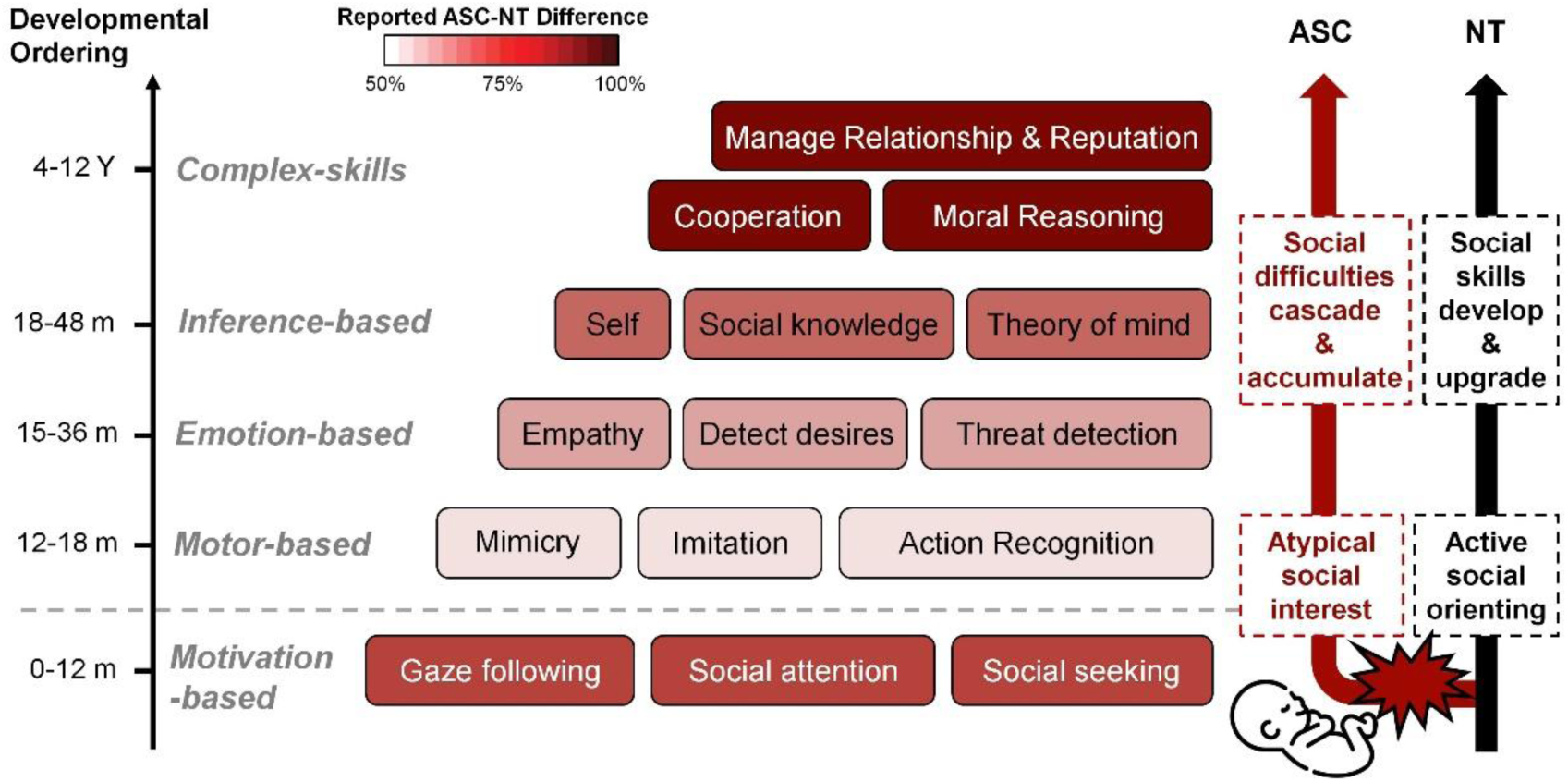
A hierarchical model of social functioning in autism. This figure illustrates social functioning as a five-level hierarchy, progressing from foundational to complex abilities. Each level represents a domain—motivation-based, motor-based, emotion-based, inference-based, and complex social skills—arranged according to prior theories of developmental sequence in NT (neurotypical) ^27^ as well as our findings of cross-sectional ordering in ASC (Fig. 4a). Darker red shading indicates greater reported differences between ASC and NT individuals (from Fig. 1c). In neurotypical development, social motivation appears in early infancy, supporting social learning through joint attention and engagement with caregivers. Motor-based skills subsequently facilitate these interactions, emotional understanding enriches them, and higher-level inference abilities refine them, together contributing to complex social functioning ^27^. In autism, early differences in social motivation may alter child-caregiver interactions, reducing opportunities for social input and interactive learning. These early differences can set off cascading effects, which are associated with progressively greater differences in higher-order social skills over time due to accumulating developmental influences. Notably, although motor-based skills arise early in this sequence, their differences in autism are less pronounced than those in motivation-based domain. This may be because imitation and action perception can develop through alternative pathways (e.g., associative sensorimotor learning and environmental feedback^60^), making them less reliant on social engagement. This hierarchical model reflects both the vertical developmental organization of social cognition and the dynamic relationships among domains, offering insight into the interconnected nature of social difficulties in autism and contributing to a broader understanding of how human social capacities develop, interact, and vary across conditions.

Our findings underscore the dynamic and malleable nature of social functioning in autism, revealing both developmental shifts and interdependent challenges across domains. Overall, social difficulties tend to intensify with age, consistent with an accumulated effect (**Fig. 4b**), where increasing social complexity in adulthood exacerbates interactional challenges. However, not all social components follow a uniform pattern of change. While foundational motor-based skills (e.g., imitation) and certain higher-order abilities (e.g., aligning behavior with social norms) improve in adulthood—likely due to learning from experience and compensatory adaptation—other skills, particularly those requiring real-time social judgment (e.g., prosocial decision-making) and flexible social adaptation (e.g., social regulation for reputation), tend to decline over time (**Fig. 4e**). These developmental changes are further illuminated by cross-domain associations. We found that autistic individuals exhibit significantly stronger correlations between emotion-based and inference-based domains compared to neurotypical peers (**Extended Data Fig. 1**), indicating tighter coupling between affective and cognitive processes in social functioning. This may reflect reduced modularity or greater interdependence between emotional understanding and mentalizing in autism^61^. Rather than operating as distinct systems, these processes may co-develop or compensate for each other, especially when intuitive social understanding is limited^62,63^. Clinically, these findings suggest that interventions targeting social difficulties in autism may benefit from integrated approaches that simultaneously engage both emotional and inferential components ^64^. For instance, emotion recognition training could be more effective when paired with explicit perspective-taking exercises, and vice versa. Addressing the interplay between emotion and inference—rather than treating them in isolation—may offer more robust pathways for enhancing adaptive social functioning ^65,66^.

Emerging perspectives, such as the Double Empathy Theory^67^ and the Social Interaction Mismatch Hypothesis^9^, challenge the traditional view that social difficulties in autism stem solely from intrinsic deficits. Instead, these theories highlight bidirectional interactional challenges, where autistic and non-autistic individuals experience mismatches in social cognition, communication styles, and expectations. Our hierarchical framework provides a structured, empirical approach to systematically assess these mismatches by quantifying 22 distinct components across five social functioning domains. This fine-grained profiling could support the development of a standardized social functioning assessment, allowing clinicians to track individualized profiles of social functioning and benchmark them against population-level norms akin to pediatric growth charts. Such profiling also enables data-driven subtyping of autism ^68^, identifying distinct social difficulties profiles rather than relying on broad diagnostic classifications. Achieving clinically meaningful subtyping will require moving beyond traditional behavioral measures—many of which our analysis indicates have limited reliability—toward multi-modal assessment batteries that integrate optimized behavioral paradigms with objective neurophysiological markers, in line with the NIH Research Domain Criteria (RDoC) framework^69^.

Beyond advancing understanding, our review identified systematic methodological issues in autism psychometrics and provides specific recommendations for improving research rigor. Sampling biases—such as underpowered sample sizes, unbalanced age distributions, and skewed male-to-female ratios (**Extended Data Fig. 2**)—undermine the generalizability and replicability of findings (see sample size recommendations in **Extended Data Table 3**). Methodological heterogeneity in tools, tasks, and measurements (**Fig. 3**) further limits comparability across studies. Importantly, however, such diversity also offers substantial advantages, enabling researchers to identify paradigms with greater ecological validity and sensitivity, and fostering methodological innovation and refinement. Earlier studies, despite variability, provided foundational insights and essential groundwork that have guided contemporary research. Building on this groundwork, our meta-analysis systematically quantifies how task sensitivity, robustness, and ecological validity influence reported effects across the autism literature. Rather than endorsing specific paradigms, we propose a data-driven framework for selecting them, emphasizing three empirically supported principles: high sensitivity to between-group differences, robustness against bias, and strong ecological validity (e.g., interaction-based, real-world assessment) ^57,70–72^ (**Fig. 3**). Applying these principles could help converge on harmonized toolkits suitable for both research and clinical assessment, enabling cross-site comparability and facilitating the translation of our five-domain framework into reliable, clinically applicable subtyping. In parallel, standardizing terminology and reporting practices, promoting data transparency and sharing, and providing detailed statistics beyond significance tests (e.g., means, standard errors, effect sizes, and behavioral correlations across multiple assessments) will be essential for improving the robustness, reproducibility, and clinical utility of autism research.

Our findings hold significant global relevance, highlighting the critical role of sociocultural factors in shaping the social difficulties experienced by individuals with autism. The significant correlation between the skewed male-to-female ratio and alterations in social functioning (**Fig. 4c**) underscores entrenched gender stereotypes in diagnostic practices. Autistic females often present with subtler or less disruptive symptoms that align more with social norms for women (i.e., the camouflaging effect), which makes them more likely to be overlooked or misdiagnosed. Consequently, those who are diagnosed often exhibit more severe symptoms ^73–76^. Future diagnostic frameworks must address this sex bias by enhancing sensitivity to subtler presentations of autism in women and ensuring more equitable diagnostic outcomes ^74,77,78^. Cultural contexts, such as low perceived social support or high levels of masculinity (reinforcing rigid gender roles and competition)—can amplify the challenges faced by autistic individuals (**Fig. 4d**). Societal pressures to conform to strict norms can amplify difficulties in social adaptation, further marginalizing individuals with autism. To overcome these barriers, public health policies should prioritize inclusive education, reduce structural inequalities, and foster more widespread acceptance of neurodiversity ^6,43^. Recognizing how cultural contexts shape the expression and recognition of autism will also be crucial for designing tailored support options and equitable care systems ^38,79^. This culturally informed approach will particularly benefit historically underserved populations while advancing evidence-based, equitable care systems.

This review, although large in scope, has several limitations. First, variability in the quality and methods of the 2,500 studies synthesized—such as differences in sample sizes, diagnostic criteria, cultural representation, and task paradigms—may have influenced our findings. The large-scale nature of this meta-analysis required grouping diverse tasks within each social component, which may obscure subtle but important differences between studies. Second, the estimation of the developmental onset of social differences could be limited by the availability of appropriate measurement tasks. The observed sequence (motivation at 6 months → motor at 12 months → emotion at 18 months → inference at 36 months) may reflect research focus and task design rather than strictly representing cognitive onset in autism. For example, while early social motivation differences are well-documented in infancy, higher-order social abilities, such as theory of mind, are rarely assessed in toddlers, making it unclear whether their later first observed differences in autism reflects true developmental timing or measurement gaps. Addressing this limitation will require the development of infant-appropriate tasks that can capture early cognitive differences. Third, the hierarchical framework of social difficulties is based primarily on cross-sectional data and requires future validation through longitudinal studies to confirm developmental patterns over time and clarify causal relationships. Fourth, this review focuses on autism without intellectual disabilities, leaving individuals with co-occurring intellectual disability underexplored. Given the significant influence of cognitive abilities on social performance and adaptation, future studies should systematically investigate how intellectual disability shapes social functioning in autism. Fifth, global representation is uneven, with limited research from low- and middle-income countries, which restricts the generalizability of our findings. Finally, while we provide recommendations, implementing these insights into standardized clinical practices and culturally sensitive policies will require interdisciplinary collaboration and longitudinal validation.

In conclusion, by offering a hierarchical framework of social functioning in autism, this meta-analysis has direct implications for clinical diagnostics, intervention development, and precision medicine approaches. Understanding the early cascading effects of social motivation alterations can inform early screening programs, while the identified cross-component compensatory patterns suggest novel rehabilitation and therapy strategies that leverage structured learning and motor-supported social engagement. These insights can be integrated into clinical decision-making algorithms, intervention trials, and digital health platforms to tailor therapies based on an individual’s specific social profile. Our findings further highlight the importance of culturally sensitive diagnostic criteria, addressing long-standing biases in autism assessment (e.g., underdiagnosis in females). Moving forward, future research should validate these insights in longitudinal and intervention-based studies to refine precision-based interventions for autism and related neurodevelopmental disorders.

## Methods

### Selection criteria and analytical strategy

In line with our conceptual framework, we included studies targeting a broad range of constructs related to social functioning—including not only observable social behaviors (e.g., communication, imitation) but also underlying cognitive, affective, and motivational processes (e.g., social attention, emotion recognition, social motivation, theory of mind)—as all contribute to an individual’s capacity for social interaction ^92^. Twenty-two key social components were identified through an extensive review of textbooks, systematic reviews, and primary literature in cognitive, developmental, and clinical psychology ^47,80,81^. Expert consultations were conducted iteratively to ensure comprehensive coverage of relevant components. See **Extended Data Table 1** for the definitions and related terms of each component. The literature search was conducted in May 2024. We identified studies published in PubMed and Web of Science between January 1, 1990, and August 10, 2025, using titles, abstracts, and keywords (for search terms, see **Supplementary Table 1**). In addition to database searches, articles were extracted from prior reviews and meta-analyses to ensure inclusiveness. Eligible studies were based on three criteria (**Fig. 2**). First, they had to be behavioral studies or had collected behavioral data in neuroimaging experiments that evaluate at least one social component. Second, they should include clinical participants diagnosed with autistic disorder, autism, ASD, high-functioning autism disorder, Asperger’s syndrome, or pervasive developmental disorder. All ASC participants were required to have a formal clinical diagnosis confirmed by qualified clinicians using standardized diagnostic instruments (e.g., DSM, ADOS, ADI, ICD, KADI, RAADS, ASDI, ASSQ, CARS) or equivalent expert-applied criteria. Studies relying solely on self-report (e.g., AQ) or ‘high-risk’ status without formal diagnosis were excluded (see **Supplementary Table 2** for full coding instructions). They should also include control groups such as neurotypically developing participants, unaffected siblings, or developmentally delayed children. Third, they had to be peer-reviewed research articles in English. Fourth, we excluded intervention studies, defined as studies applying controlled manipulations (e.g., behavioral training, pharmacological treatment, neurostimulation, educational programs) to autistic participants and assessing effects on outcomes over time. These were excluded to ensure that our synthesis captured naturalistic patterns of social functioning, uncontaminated by treatment effects, consistent with prior systematic reviews^50,82,83^.

A total of 2,622 studies met our broad inclusion criteria and were included in the qualitative systematic review. These studies assessed group differences in at least one component of social functioning in autistic and neurotypical participants. For the quantitative meta-analysis, we applied stricter eligibility criteria: a) Task comparability —only paradigms demonstrating sufficient methodological homogeneity were retained for cross-study effect size synthesis; and b) Statistical completeness—studies were included only if they reported sufficient data (e.g., means and standard deviations, or t/F-values with exact p-values) to compute standardized effect sizes. Applying these criteria yielded a final dataset of 1,388 independent effect sizes for quantitative meta-analysis (with 34 tasks across 13 components) (see the PRISMA flowchart in **Fig. 2**).

This study involved two complementary meta-analytic approaches: (1) Group-difference meta-analysis—we synthesized standardized mean differences between autistic and neurotypical participants using Hedges’ *g* as the effect size metric. These models estimated the magnitude of group differences for each task and social functioning component, and assessed potential moderators (e.g., age, IQ, measurement type). (2) Cross-domain correlation meta-analysis—to quantify relationships between social functioning domains. Pearson’s *r* values were converted to Fisher’s Z prior to synthesis to stabilize variance estimates, and then back-transformed for reporting purposes.

### Data extraction and coding quality assessment

Following a Codebook (**Supplementary Table 2**), twenty well-trained postgraduate students extracted information from included articles. Information includes publication details (article name, journal, year of publication, study type, impact factor, citation counts), participant demographics (participant type, clinical population, diagnostic criteria, nationality, age group, mean age, sample size, mean IQ, clinical number, female included, male included, control population, mean age of the control group, diagnosed comorbidity), methodological details (task paradigm, index, task design), and outcome details (conclusion, effect size, and behavioral correlations with other tasks). The ecological validity of each task was rated ^70^, following the criteria outlined in **Supplementary Table 3**. We also coded the reporter type (self vs. proxy) for survey-based measures, task index (accuracy vs. reaction time) for action perception studies, and measurement modality (task-based vs. survey-based) for the entire literature.

To enhance coding reliability, each human coder was assigned to only 2∼4 components to ensure they had a good understanding of the studies and were familiar with the coding of component-specific variables. Ten percent of studies were coded by two or three independent coders, and the inter-coder correlation (Pearson’s *r*) was 0.91 (comparable to other large-scale meta-analysis projects^84^). Discrepancies were resolved through consensus meetings and consultation with senior researchers.

According to the author’s conclusion (p-value of group main effect between the autistic and non-autistic group), each study was coded into one of five categories: impaired, different, complex, intact, and outperformed. Operational definitions for each category are detailed in **Supplementary Table 2** to ensure coding consistency across studies. *Impaired* means the autistic group performed significantly poorer than the non-autistic group, *intact* means no significant differences, *outperformed* means that the autistic group outperformed the non-autistic group, *different* means that the autistic group performed differently from the non-autistic group and did not indicate inferior or superior performance. When the conclusion varied across different experimental conditions, it would be marked as *complex*.

This five-category system was used to support a qualitative synthesis of global trends in reported outcomes, especially for visualizing the overall distribution of alterations across 22 social components (**Fig. 1c**). Many studies could not be included in quantitative meta-analysis due to incomplete statistical reporting (e.g., missing means or standard deviations) or highly heterogeneous experimental designs, making effect size estimation infeasible. The five-category coding thus enabled the inclusion of a broader evidence base, maximizing the scope of our review. Crucially, this categorization was not used in any of the main quantitative analyses. All effect size calculations, moderator tests, and behavioral correlation analyses were conducted using continuous metrics (Hedges’ *g*).

To address the ambiguity and arbitrariness in terminology usage across the literature, we adhered to a standardized coding strategy. Specifically, for tasks with inconsistent component labels (e.g., gaze preference tasks have been labelled in the literature either as social attention or social motivation; read mind in the eyes (RMET) have been labelled either as theory of mind or emotion perception), we adopted the label most commonly used in the field, rather than relying on the terminology arbitrarily applied by individual study authors.

### Categorize social difficulties in autism into five domains

Four prominent neurocognitive theories of autism (each citation > 2,000) guided our grouping of social difficulties in autism. The broken mirror theory builds on alterations in motor-based components like action perception and imitation ^85,86^. The empathizing-systemizing theory is associated with emotion-based components like emotion perception and empathy ^87^. The mind-blindness theory ascribes autism to components involving understanding self and others, such as theory of mind and social reasoning ^88^. The social motivation theory centered on components like social attention and reward responsiveness ^29^. Components not explicitly covered by these theories, such as social learning, pretend play, and communication, were grouped as complex skills, reflecting higher-order social abilities. This systematic approach ensured logical, evidence-based grouping within established autism research frameworks.

### Statistical Analysis of Effect Sizes

To ensure the stability and robustness of our meta-analytic estimates, only task paradigms with data from at least four independent studies were included in the quantitative synthesis of effect sizes. To standardize outcomes across studies, Hedges’ *g* was selected as the primary effect size metric due to its correction for small-sample bias. For components where multiple outcome indices were available, we selected the most frequently reported index across the literature for that component to ensure consistency in our analysis. Effect sizes were calculated from reported statistics (e.g., means, standard deviations, t-values, or F-values) using the *esc* package^89^ in R. By convention, a negative Hedges’ *g* indicates that the ASC group performed more poorly than the neurotypical group. This process yielded a standardized effect size and its corresponding variance for every eligible outcome, preparing the data for meta-analytic synthesis. To enhance transparency and interpretability, forest plots for each component were provided in OSF repository (https://osf.io/jzw45/files/osfstorage).

All meta-analyses were conducted using random-effects models in conjunction with robust variance estimation (RVE). This approach was necessary because many included studies contributed multiple, potentially correlated effect sizes, including multiple outcomes reported within the same task and social component (e.g., separate stimulus categories, experimental conditions, subscales, or participant subgroups). These finer-grained outcomes were reported inconsistently across studies, and the correlations among them were rarely available. As a result, the within-study dependence structure was heterogeneous and only partially observed, making it difficult to specify a single multilevel or cross-classified random-effects structure. To robustly account for this non-independence without imposing strong structural assumptions, we employed RVE as implemented in the *robumeta* package in R, clustering effect sizes by study and specifying the correlated-effects (CE) working model. The intra-study correlation (ρ) was set to 0.5 ^90,91^ and verified through sensitivity analyses showing that varying ρ between 0 and 0.9 did not meaningfully affect the results. Heterogeneity was quantified using the I² statistic and the variance of the true effect sizes (τ²).

Moderation analysis included two levels: study level and country level (for the definition of each factor, see **Extended Data Table 2**). Exploratory meta-regression was employed to test the existence of moderating effects (using the R package *robumeta*). For categorical factors, we compared the effect sizes of all categories using the Wald F test. For continuous moderators, we assessed their association with effect sizes via meta-regression coefficients tested with two-tailed t-tests. Countries with less than five studies were excluded from the meta-regression of sociocultural factors to ensure model stability. Here no adjustments were made for multiple comparisons, as these moderator analyses were exploratory and intended to identify potentially relevant demographic and sociocultural factors rather than to provide definitive hypothesis tests.

### Meta-Analysis of Behavioral Correlations

To characterize how different domains of social functioning relate to one another, we conducted the correlation analyses in two stages, separating the comparison of association strength from the evaluation of higher-order organizational structure. We extracted all available behavioral correlation coefficients across the five social functioning domains. Correlations were categorized by domain pair (e.g., emotion–inference) and converted to Fisher’s z scores prior to analysis.

In the first (exploratory) stage, we compared the strength of cross-domain correlations between autistic (ASC) and neurotypical (NT) groups using a single, unified correlated-effects robust variance estimation (RVE) meta-regression model. All Fisher’s z-transformed correlations from both groups and all domain pairs were analyzed simultaneously within this model, with domain pair and diagnostic group (ASC vs NT) treated as categorical predictors. Effect sizes were clustered by study to account for dependency arising from multiple correlations contributed by the same study. This approach allows pooled correlations for each domain pair and group to be estimated concurrently, rather than via separate subgroup meta-analyses. Group differences for each domain pair were tested using Wald-type linear contrasts within the same model, with small-sample corrected variance estimates (CR2) and false discovery rate (FDR) correction for multiple comparisons (**Extended Data Fig. 1b**).

In the second (confirmatory) stage, we examined directional dependencies among domains using meta-analytic structural equation modeling (MASEM). For each study, we constructed a 5 × 5 correlation matrix; when multiple correlations were available for a given domain pair, they were aggregated into a single variance-weighted estimate. Using the *metaSEM* package, we compared three theory-driven directional models (**Extended Data Fig. 1c**): (1) a Serial model, where influence flows sequentially from motivation to complex skills following the developmental order identified in **Fig. 4a**; (2) a Radial model, in which motivation-based processes directly influence all other domains; and (3) a Network model allowing all pairwise connections. Model fit was evaluated using the Bayesian Information Criterion (BIC), and group differences in path strengths were tested using likelihood ratio tests. The robustness of the best-fitting model was further evaluated via permutation testing against 120 alternative unidirectional cascade permutations.

### Publication Bias Assessment

For each task paradigm with ≥10 studies, we tested for publication bias using an RVE-adjusted approach. Specifically, we applied Egger’s regression within a robust variance estimation framework (robumeta), clustering effect sizes by study to account for within-study dependencies. A significance threshold of p < 0.10 was used for detecting funnel plot asymmetry. When asymmetry was present, PET-PEESE corrections were conducted within the same RVE model to obtain adjusted effect size estimates. This method preserves statistical power and precision while properly accommodating non-independent data structures.

### Pre-registration and Deviations

This meta-analysis was pre-registered on PROSPERO in July 2024 (www.crd.york.ac.uk/PROSPERO/view/CRD42024566141). The pre-registration outlined key inclusion criteria (e.g., participants aged ≥3 years, IQ ≥75), search strategies, coding procedures, and statistical models (e.g., random-effects meta-analysis, meta-regression). During study screening and coding, two deviations from the protocol were made. First, we expanded the age inclusion range to allow studies including infants under 3 years, provided they were confirmed to have ASC through longitudinal follow-up or validated early diagnostic assessments. This change was based on growing evidence for the predictive value of early social markers and impacted 94 studies (3.59% of total). Second, in cases where individual IQ data were unavailable, we included studies only if the mean IQ of ASC group was ≥75 or if the authors reported that their participants did not have intellectual disability. These modifications were documented prior to analysis and did not qualitatively alter the main findings (see sensitivity tests in **Supplementary Fig. 3**).

Following editorial guidance during the final revision stage, we conducted an additional updated literature search to ensure that the meta-analysis reflected the most recent research in the field. The original pre-registration covered studies published up to March 2024. The updated search, using the same databases and search strategy, extended the coverage to all eligible studies published before August 2025. This update identified 3,886 additional records, which after screening resulted in 141 additional studies included in qualitative synthesis and 40 additional studies included in quantitative meta-analyses, and expanded the geographical coverage from 29 to 32 countries. All newly included studies were coded using the same procedures and criteria as the original dataset. Re-running the analyses with the expanded dataset produced only minor numerical changes in some statistics, while all key findings and statistical significance patterns remained unchanged, indicating that the conclusions of the study are robust to the inclusion of the most recent literature.

## Data Availability

All data in this project have been deposited in the Open Science Framework (https://osf.io/jzw45).

## Code Availability

All data analysis code in is available on the Open Science Framework repository(https://osf.io/jzw45).

## Acknowledgements

We thank Jialu Yu, Jiamin Bao, Xiuli Zheng, Liangyu Liu, Yi Li, Mingxue Fu, Hengsen Dai, Yuxing Yang, Xiaoxiao Zeng, Lingfang Wang, Xinru Liu, Yutong Zhao, Yuchen Liu, Nan Zhang, and Bingsu Wang for literature coding and manual check, Shuxian Jin for valuable discussions and comments. This work was supported by the National Natural Science Foundation of China (32422033, 32430041 to Y.W.; 32371139, 32000776 to Y.Z.), the National Science and Technology Innovation 2030 Major Program (2022ZD0211000, 2021ZD0200500 to Y.W.), the Fundamental Research Funds for the Central Universities, and the Open Research Fund of the State Key Laboratory of Cognitive Neuroscience and Learning (Grant No. CNLZD2103 to Y.Z.).

## Author Contributions Statement

S.L., H.W., designed and planned the research. S.L., H.W., G.C., X.C, Y.W., W.H. performed the literature search, data coding and data analysis. S.L., H.W., G.C. wrote the initial draft of the manuscript, and A.H., L.S., L.Y., K.A., Z.Z., X.Z. edited and reviewed the final manuscript. Y.W. and Y.Z. conceived and supervised the project and were involved in all aspects of the research.

## Extend Data

**Extended Data Fig. 1.**
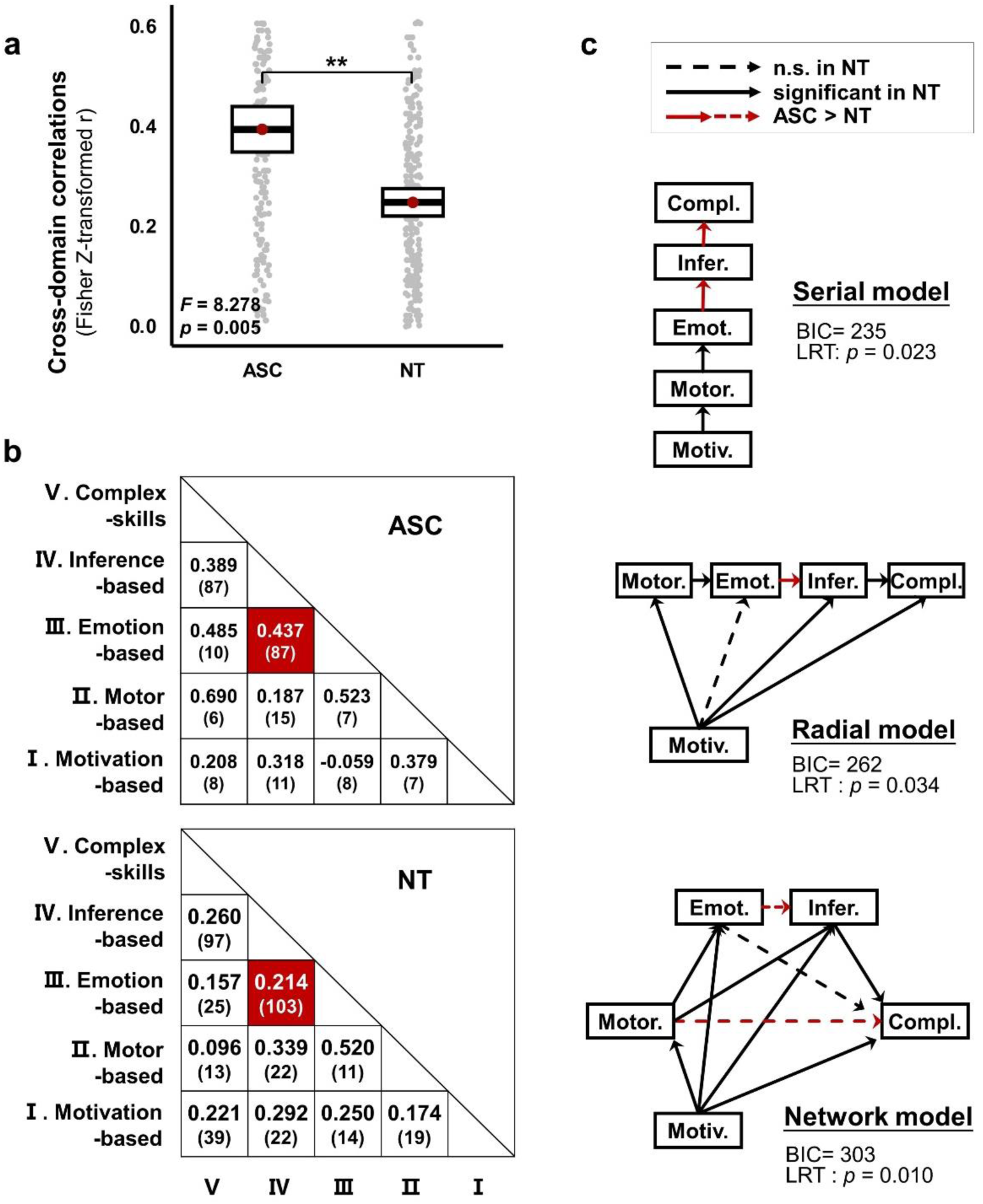
Cross-domain correlations and meta-analytic structural equation modeling (MASEM) of social functioning domains. **(a)**ASC individuals showed stronger cross-domain behavioral correlations across the five social domains than NT individuals (*F* (1, 98.2) = 8.278, partial R^2^ = 0.078, 95% CI = [0.008, 0.201], *p* = 0.005). The center line of the box plot represents the mean estimate of cross-domain correlations(Z(ASC) = 0.392, Z(NT) = 0.246), the gray dots represents individual study-level effect sizes (Fisher’s Z-transformed correlations), and the bounds of the box demonstrates the ±1 standard error(SE). **(b)** Cross-domain correlation matrices for ASC (upper triangle) and NT (lower triangle), with study counts in parentheses. Significant ASC > NT differences are in red, strongest between emotion and inference (r = 0.437 vs. 0.214, post hoc Cohen’s q = 0.252, 95% CI = [0.091, 0.413], *F* = 10.385, *p* = 0.034). Group differences for each domain pair were tested using Wald-type linear contrasts with small-sample corrected variance estimates (CR2), and p-values were adjusted for multiple comparisons using the false discovery rate (FDR) correction. **(c)** MASEM model comparison. Three theoretically derived directional architectures were compared: Serial model (influences proceed along the developmental sequence: motivation → motor → emotion → inference → complex skills), Radial model (social motivation directly influences all other domains), and Network model (all pairwise influences allowed). Paths were constrained to follow the developmental order proposed in prior theory as well as found in Fig. 4**a**^12,27^. The Serial model provided the best fit (lowest BIC) and outperformed 120 alternative unidirectional permutations (permutation test, 3/120, *p* = 0.025). Likelihood ratio tests (LRT) revealed significant group differences in specific paths (*p* = 0.024), with ASC individuals showing stronger influences from emotion to inference and from inference to complex skills (red arrows). Dashed arrows indicate non-significant paths in NT. No adjustments were made for multiple comparisons. All statistical tests were two sided.

**Extended Data Fig. 2.**
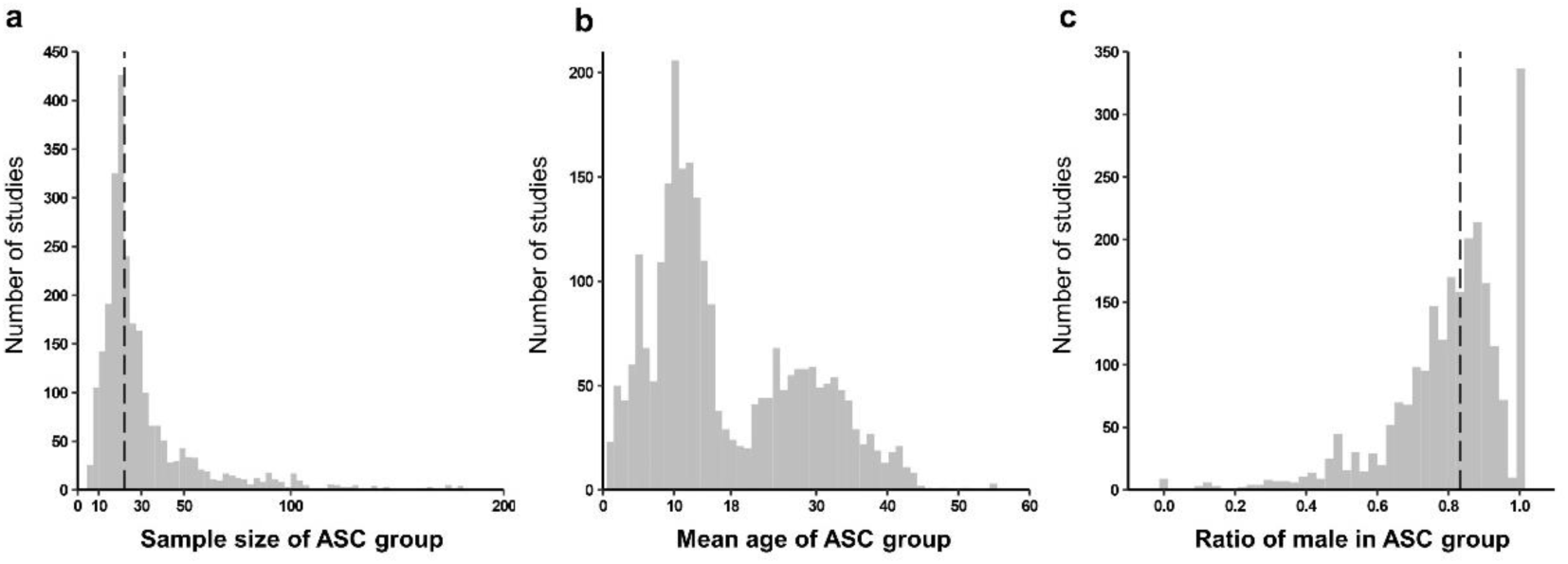
Biased distribution of studies (a) across sample size, (b) mean age and (c) ratio of male in ASC group. The median sample size was 22, and that of male ratio was 0.833 (indicated by dash lines). The mean age showed a bimodal distribution: childhood (<18 yrs) and adulthood (>18 yrs).

**Extended Data Fig. 3.**
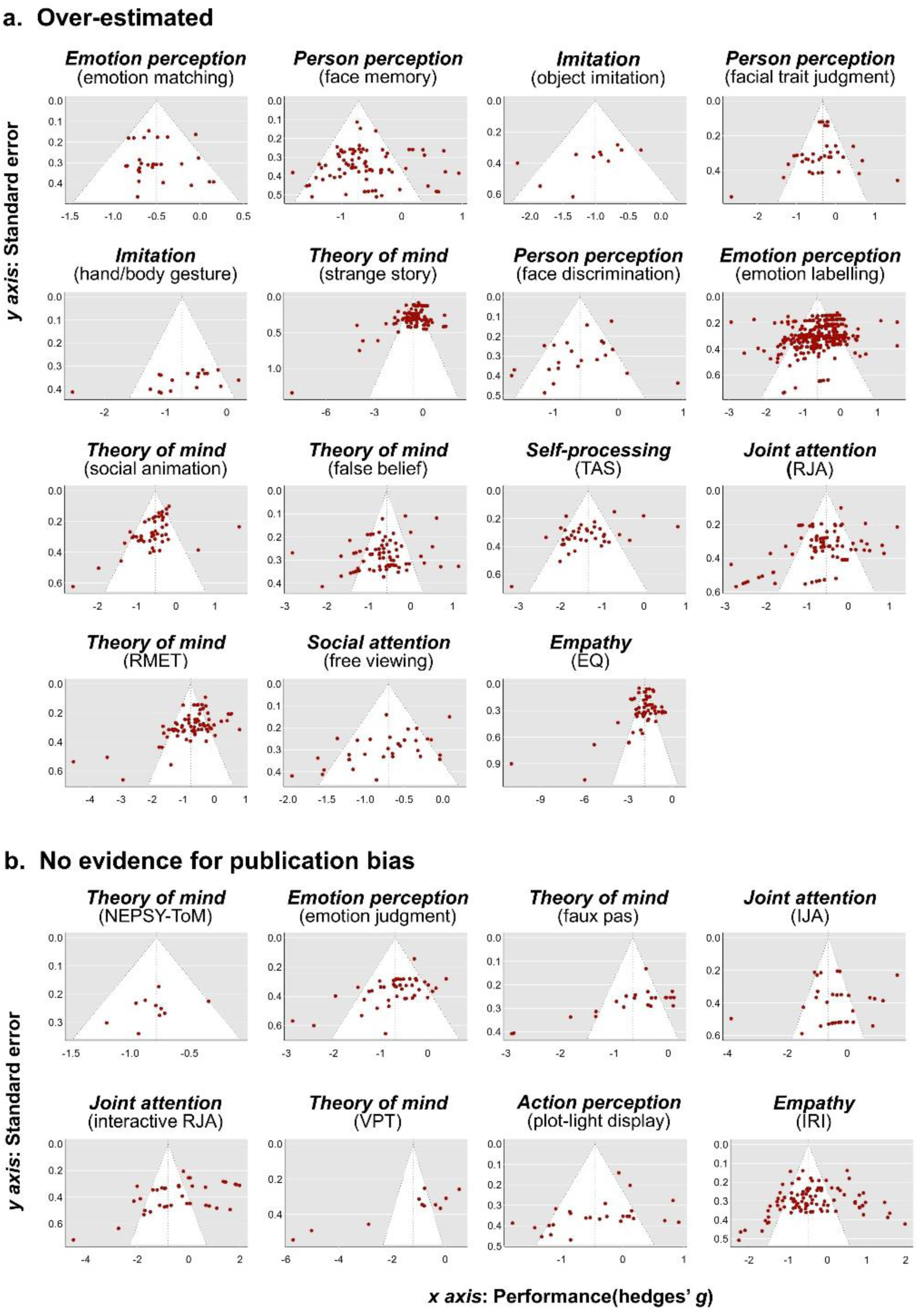
Funnel plot for each task. **(a)** ‘Over-estimated’ indicated the existence of publication bias. **(b)** ‘no evidence’ indicated the egger’s test was not significant, X-axis represents the Hedges’ *g* value (ASC-NT difference), and the y-axis represents the standard error. Negative values on the x-axis indicate studies that reported poorer performance in individuals with autism. The symmetrical triangular region represents the expected distribution of studies under no publication bias, with points distributed around the centerline (zero effect size).

**Extended Data Fig. 4.**
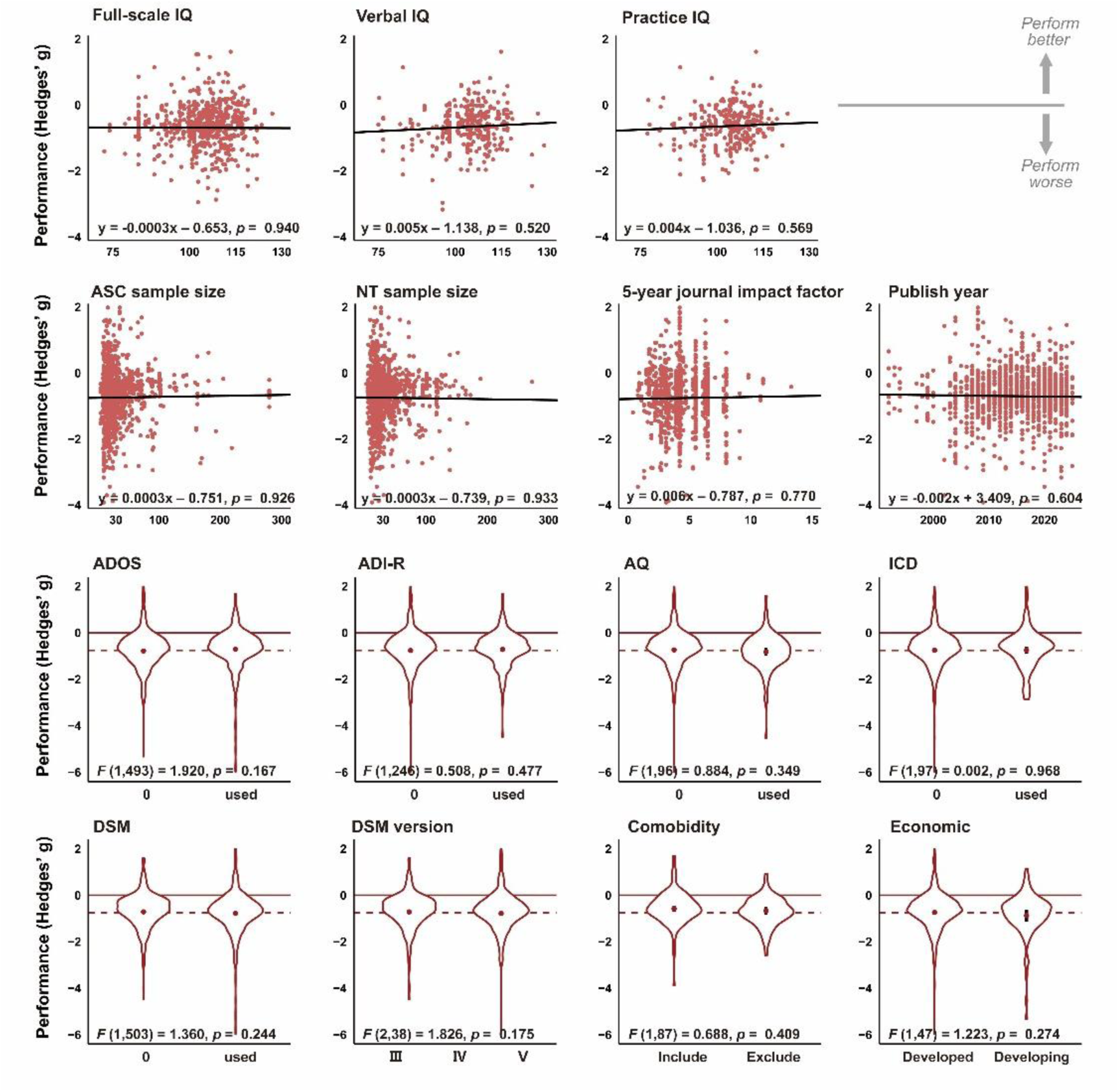
Non-significant moderators. Study-level moderators include cognitive measures (Full-scale IQ, Verbal IQ, Performance IQ), sample sizes for ASC and NT groups, publication metrics (5-year journal impact factor, publication year), diagnostic criteria (ADOS, ADI-R, AQ, ICD, and DSM versions III, IV, and V, “0” indicates studies that did not use the specified tool as a diagnostic criterion), comorbidity inclusion, and economic status (developed vs. developing countries). Data are represented as scatter plots with regression lines or violin plots for categorical variables; Regression lines and *p*-values indicate the significance of correlations after hyperbolic and symmetric Gaussian transformation. Hedges’ *g* values are plotted on the y-axis, indicating altered performance (i.e., ASC-NT difference, more negative values indicate poorer performance in ASC). Red dots and error bars in violin plots indicate the estimated effect sizes and 95% CI, respectively; All statistical tests were two sided, and no adjustments were made for multiple comparisons.

**Extended Data Table 1.**
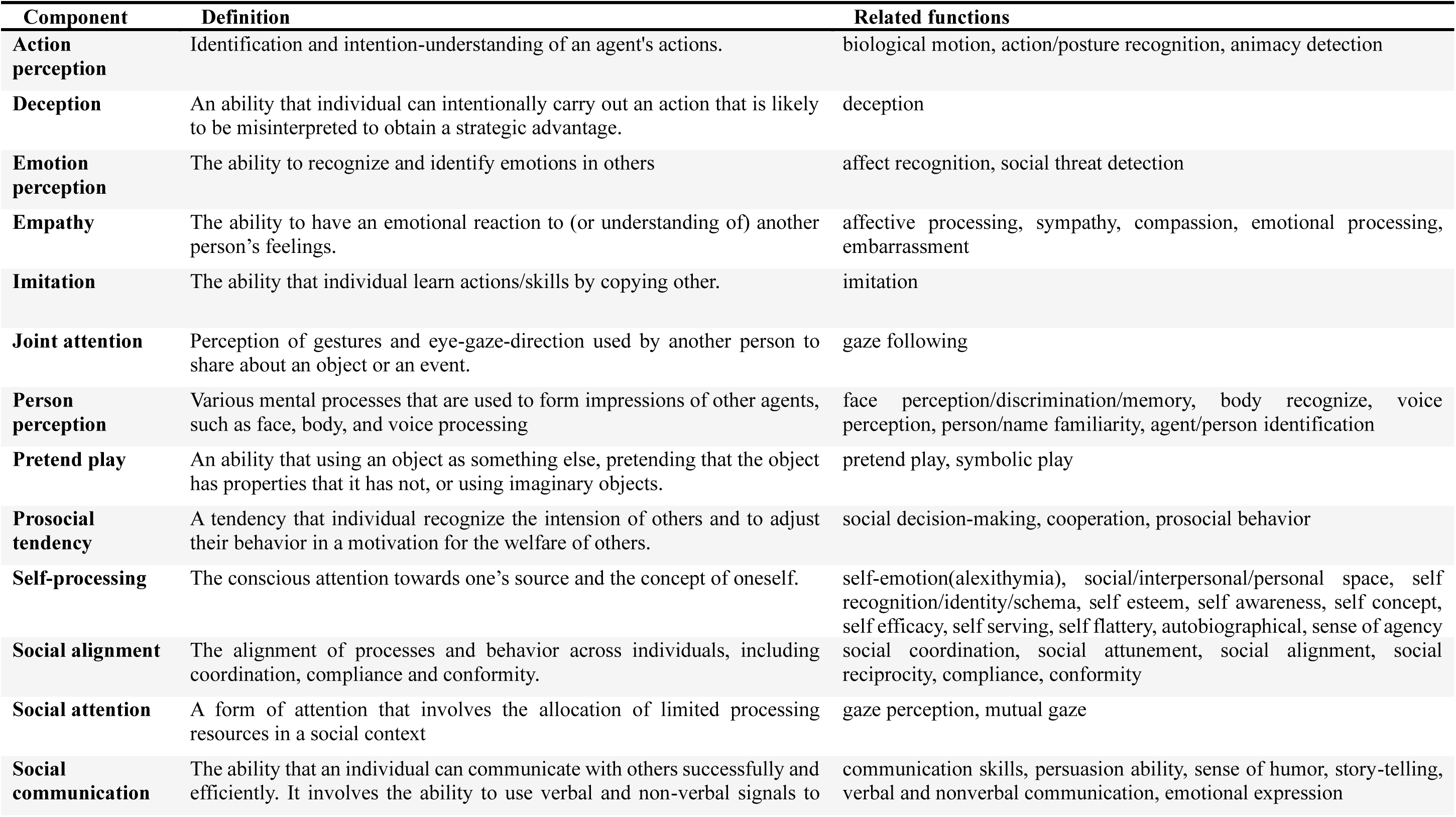
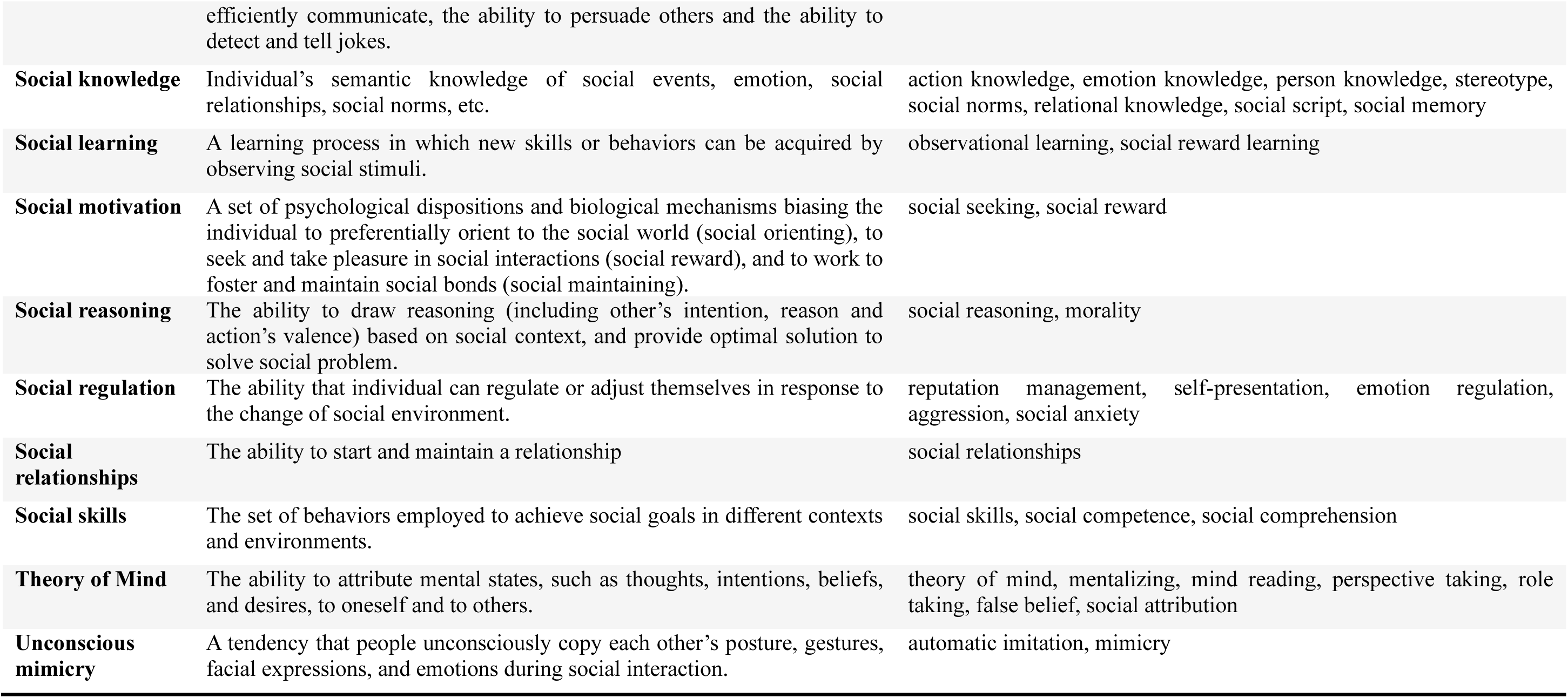
List of 22 social components, definitions, and related functions.

**Extended Data Table 2.**
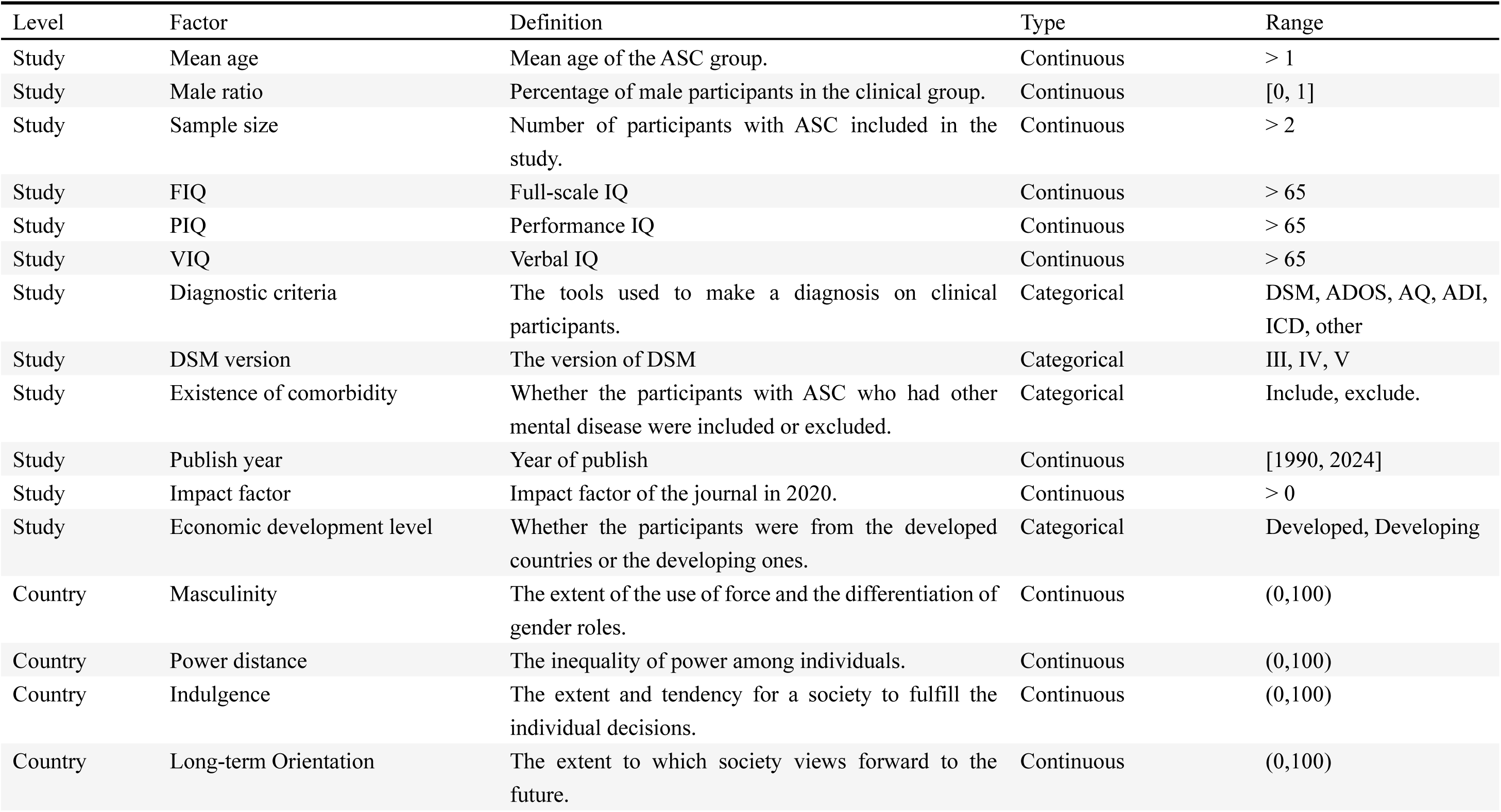
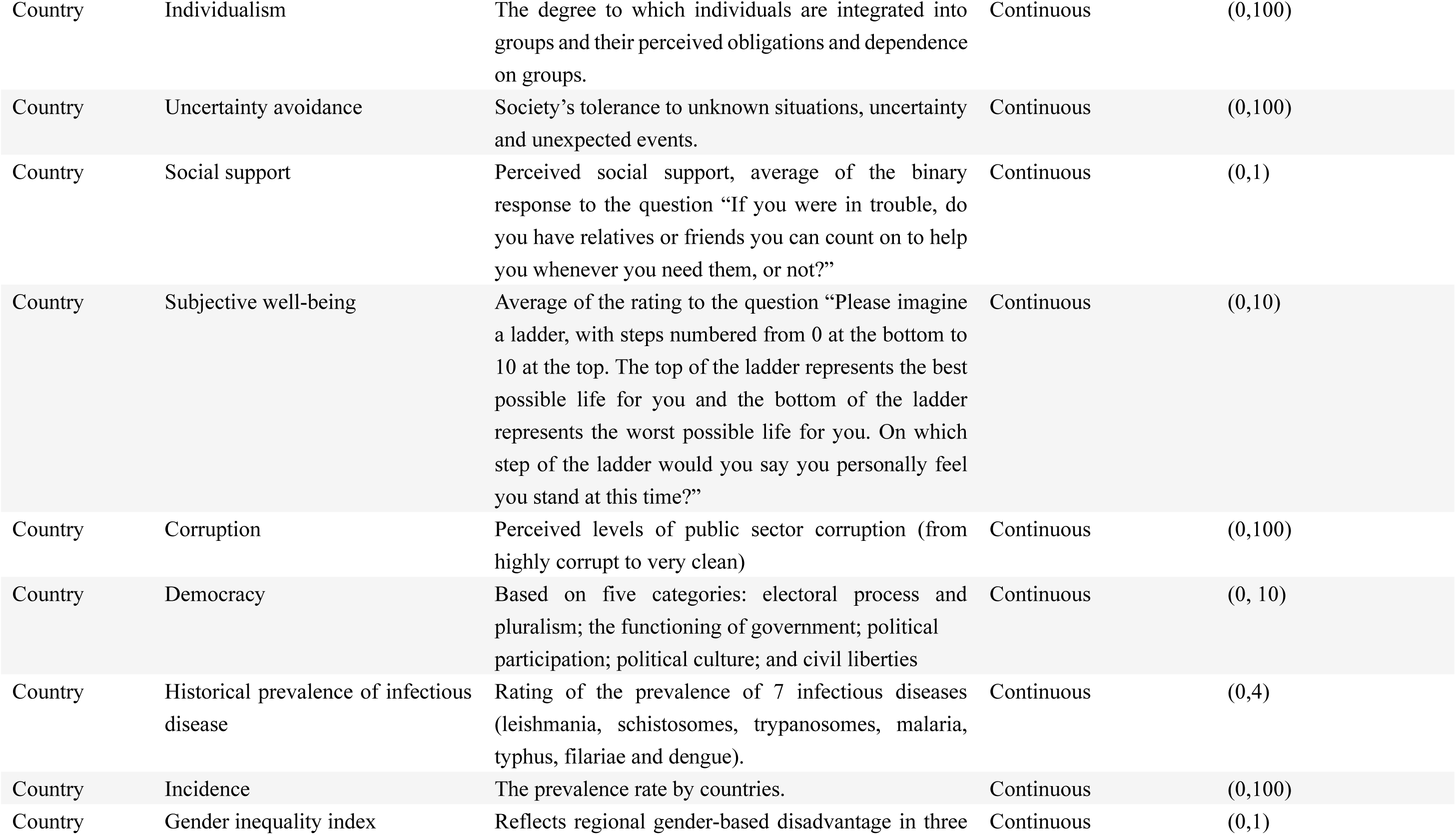
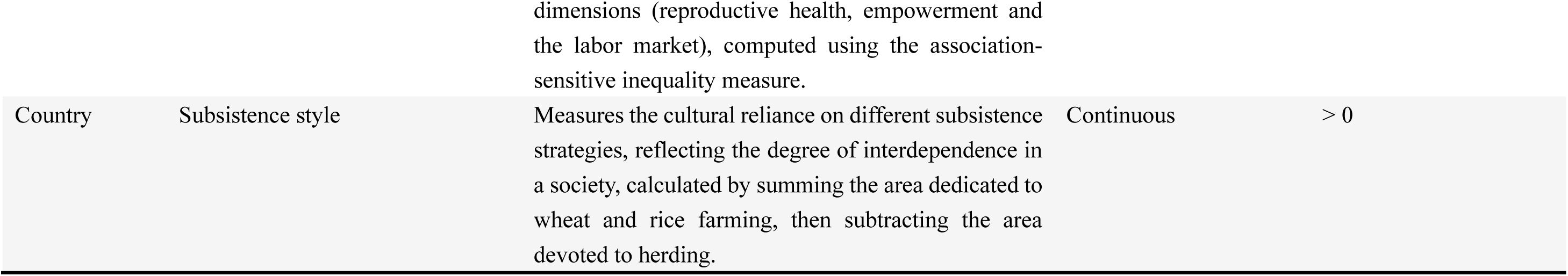
Definitions of 25 moderating factors. There are 2 levels, study level means that the values of factors were coded by each study, country level means that the effect size was grouped by the countries before meta-regression, countries with less than five studies were excluded to control the bias.

**Extended Data Table 3.**
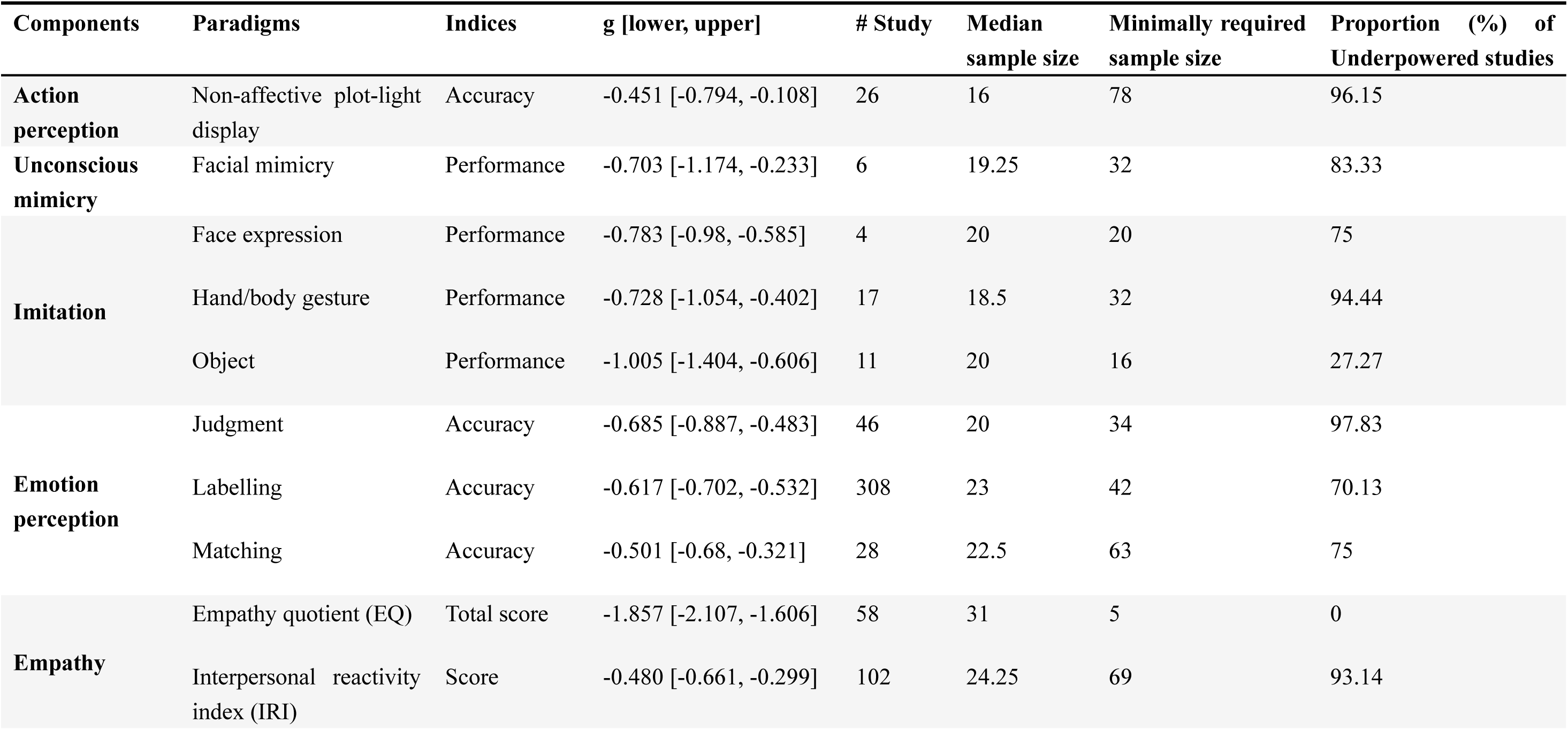
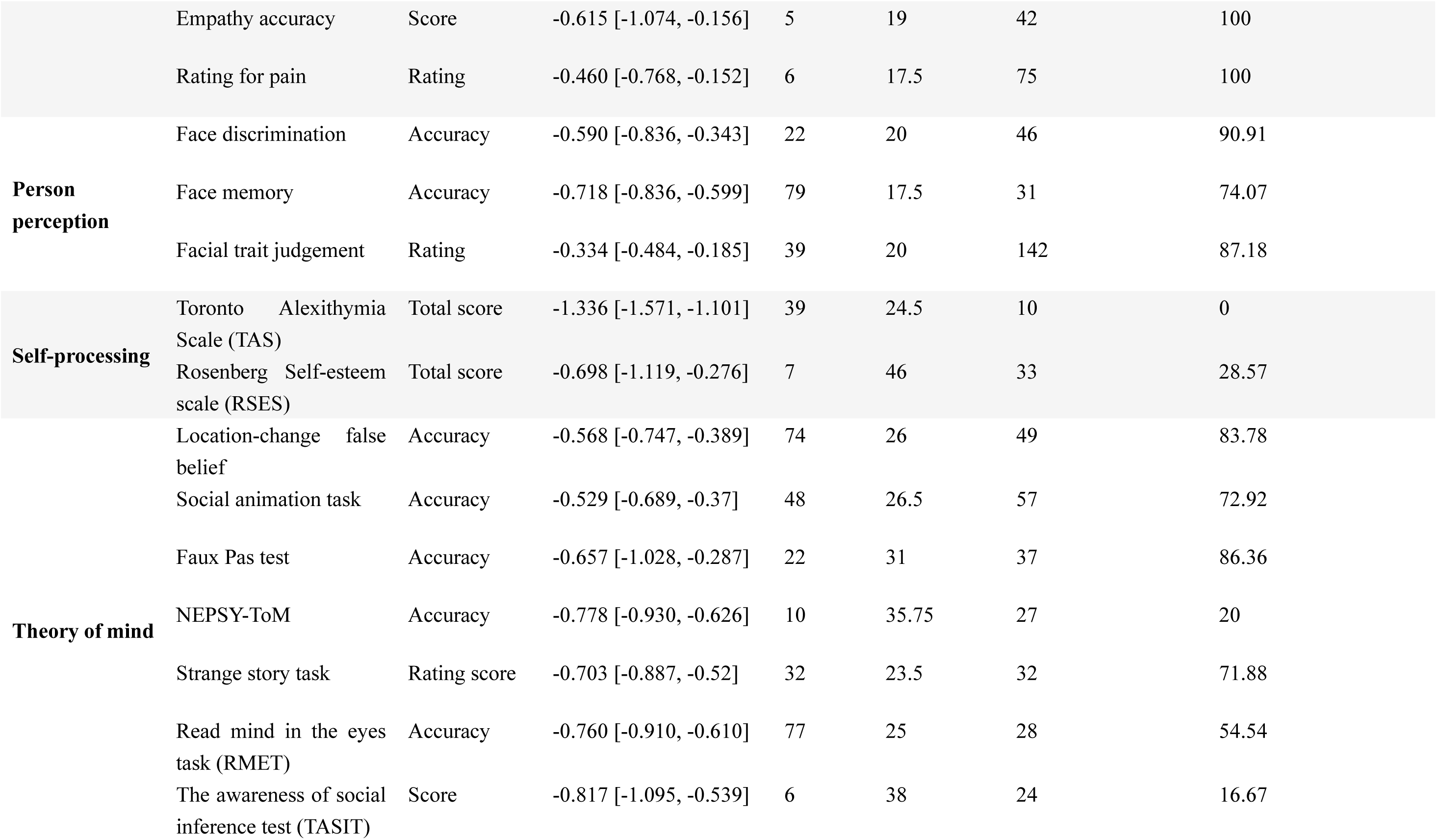
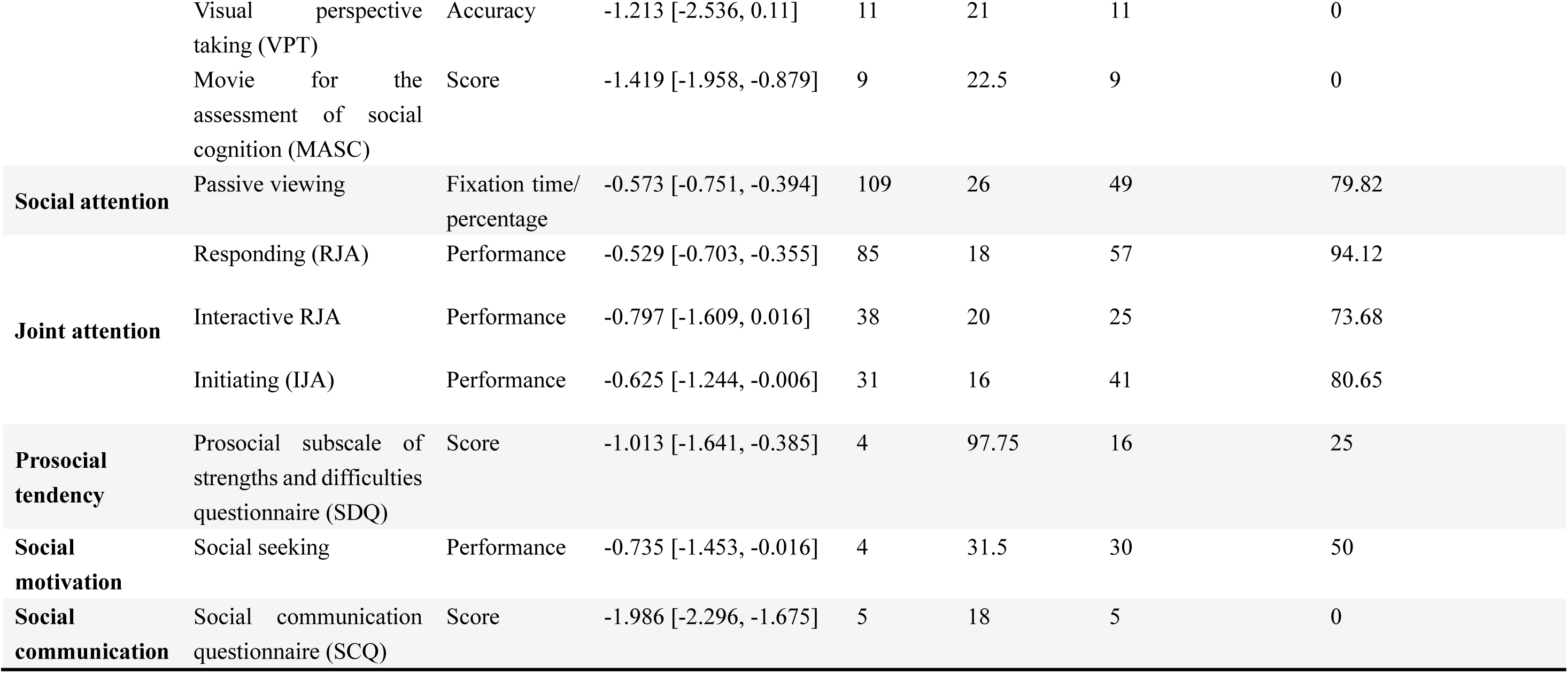
Power analysis with recommendations of sample size for each task paradigm. Columns include components, paradigms, indices, effect size (Hedges’ *g*) and range of random effect, study number (# study), median sample size, minimally required sample size (based on Power Analysis Estimation, power of 0.8, two-sided test), and proportion of underpowered studies (with sample size below the minimally required one). Tasks containing less than 4 studies were not included. We recommend future research to recruit autistic participants larger than the minimally required sample size.

## Supplementary Material

**Supplementary Fig. 1.**
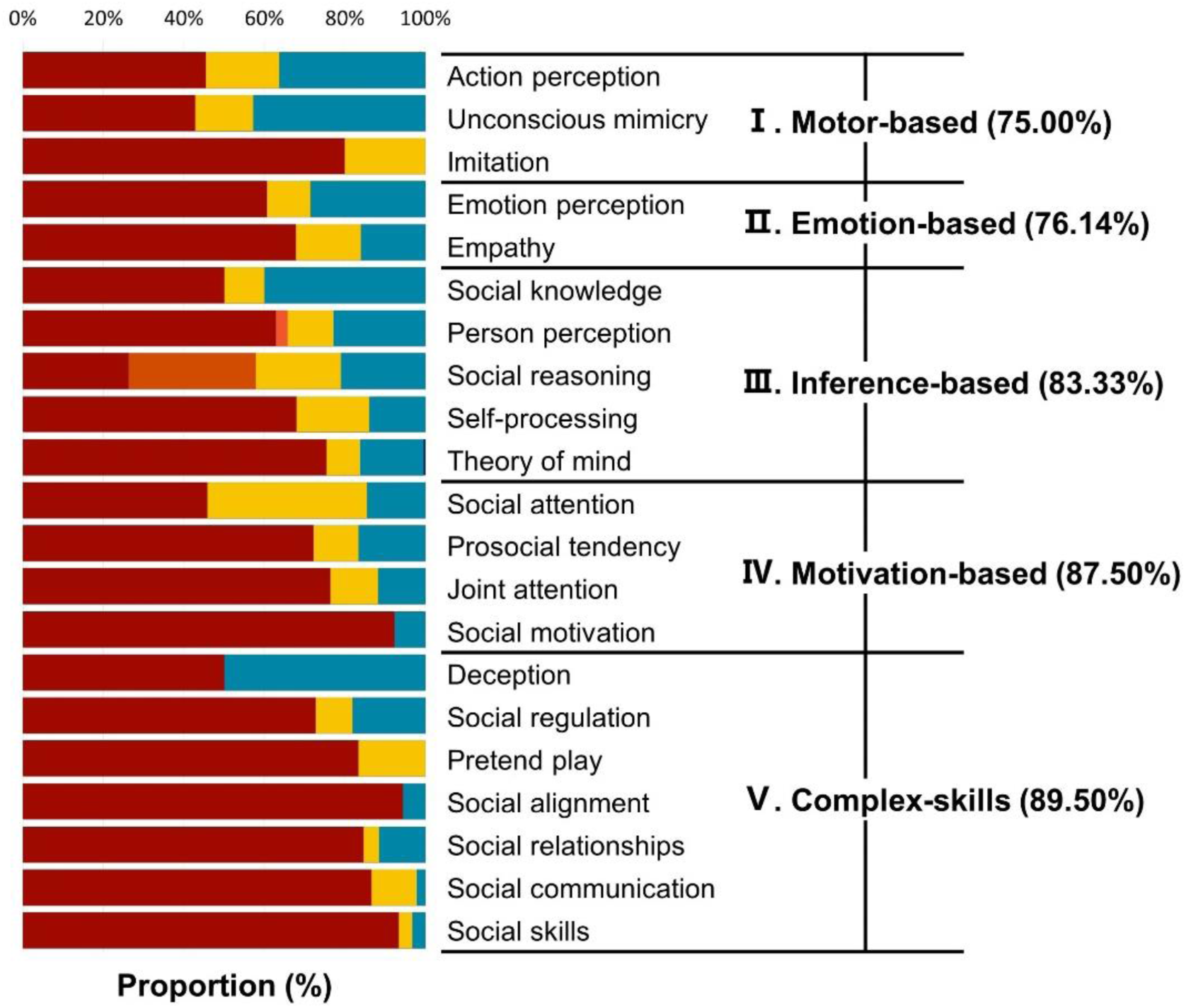
Sensitivity tests of excluding small sample size studies (N < 30).

**Supplementary Fig. 2.**
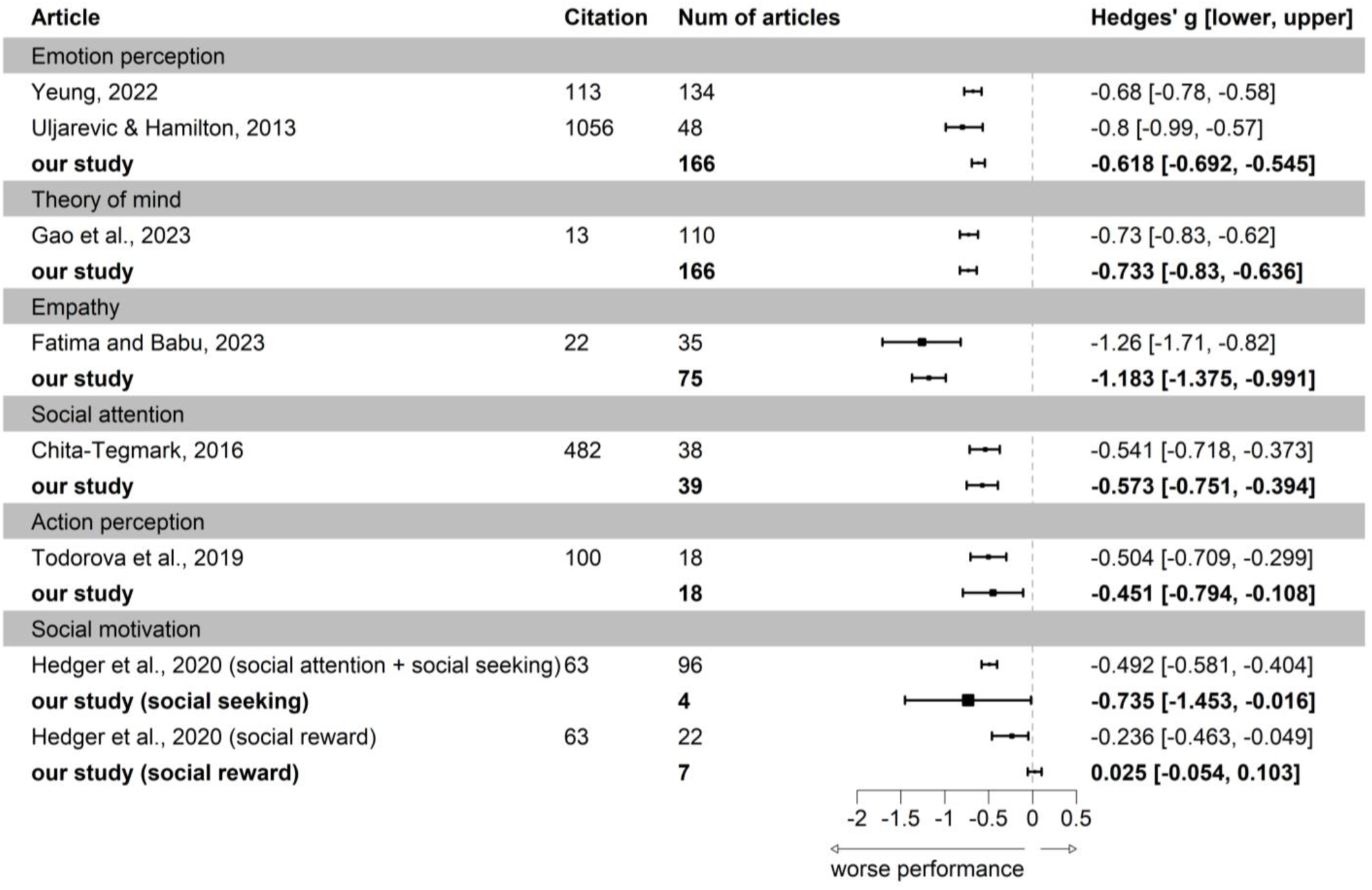
Testing the reliability of our study by comparing our results with previous meta-analyses. For each component, we only selected its most recent meta-analyses or the one with the highest citations (as of February 1, 2025). Effect sizes previously reported as Cohen’s d were converted to Hedges’ *g* with 95% confidence intervals. Negative effect sizes indicate worse performance for autism compared with neurotypical peers. Though our meta-analysis is the latest and includes much larger number of studies, our effect sizes are highly consistent with prior meta-analyses, supporting the robustness and reliability of our findings^1–7^

**Supplementary Fig. 3.**
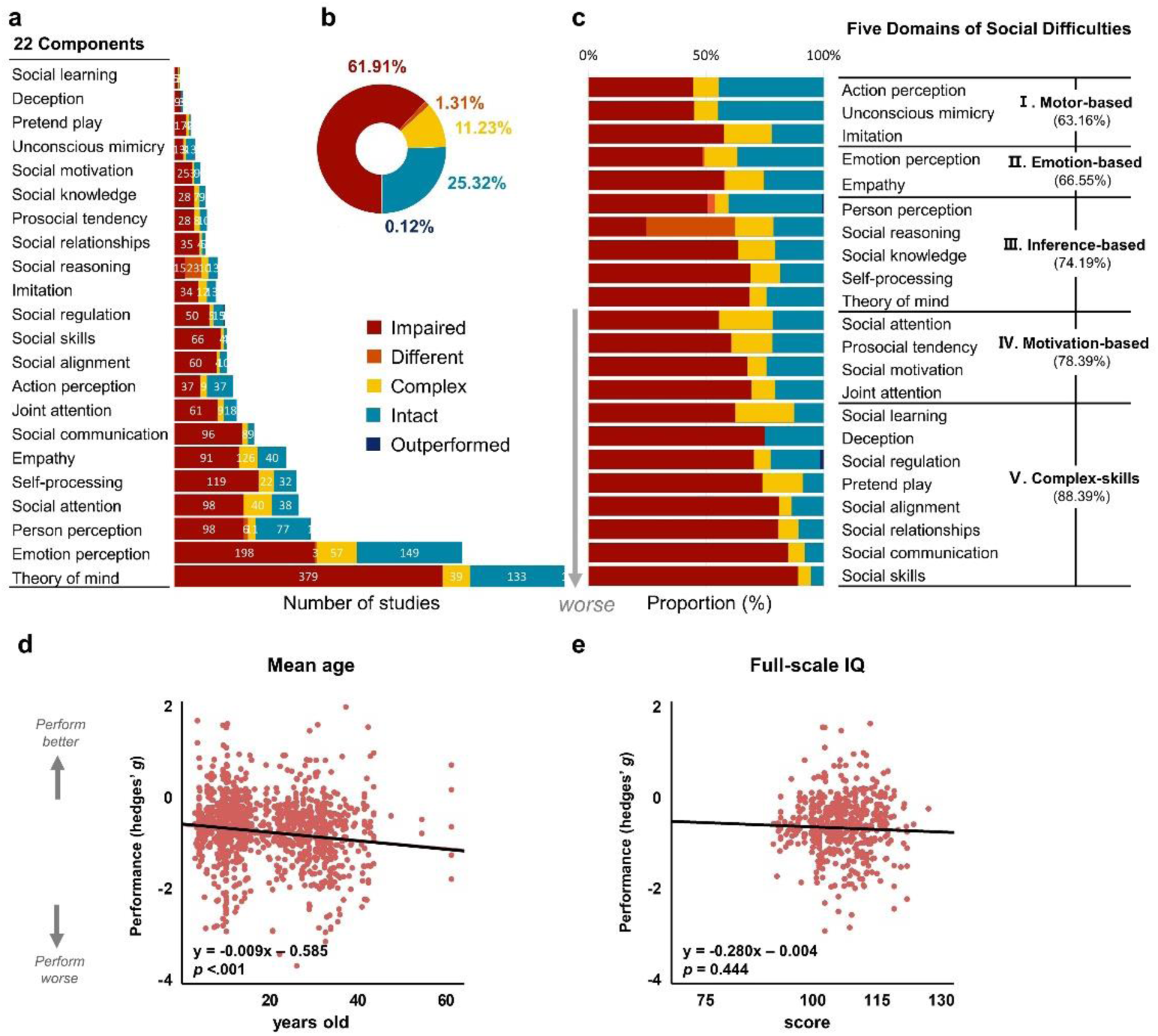
Deviation from pre-registration and sensitivity test. **(a-d)** Study distribution, proportion of conditions across all studies, prevalence of alterations in each component, and the moderating effect of age (i.e., mean age of autistic samples) all remained stable after excluding infants (age under 3) diagnosed with ASC. **(e)** The moderating effect of full-scale IQ remains stable after excluding studies containing ambiguous IQ reporting (e.g., with no individual IQ information, or IQ range cover <75 so that certain proportion of participants’ IQ under 75). Hedges’ *g* values on the y-axis represent altered performance (i.e., ASC-NT difference, more negative values indicate poorer performance in ASC). Regression lines and *p*-values indicate the significance of correlations after hyperbolic and symmetric Gaussian transformation. All statistical tests were two sided, and no adjustments were made for multiple comparisons.

**Supplementary Table 1.**
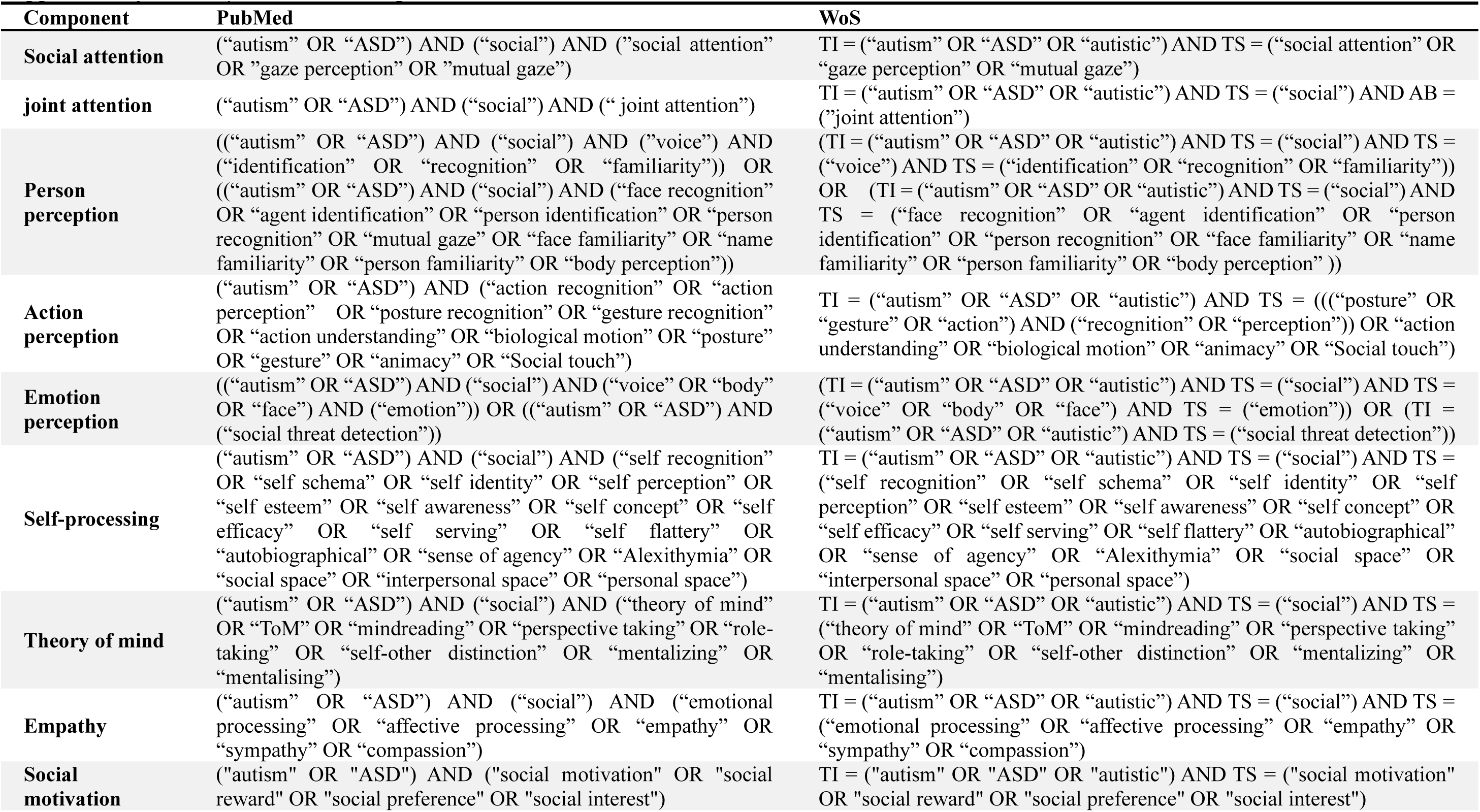
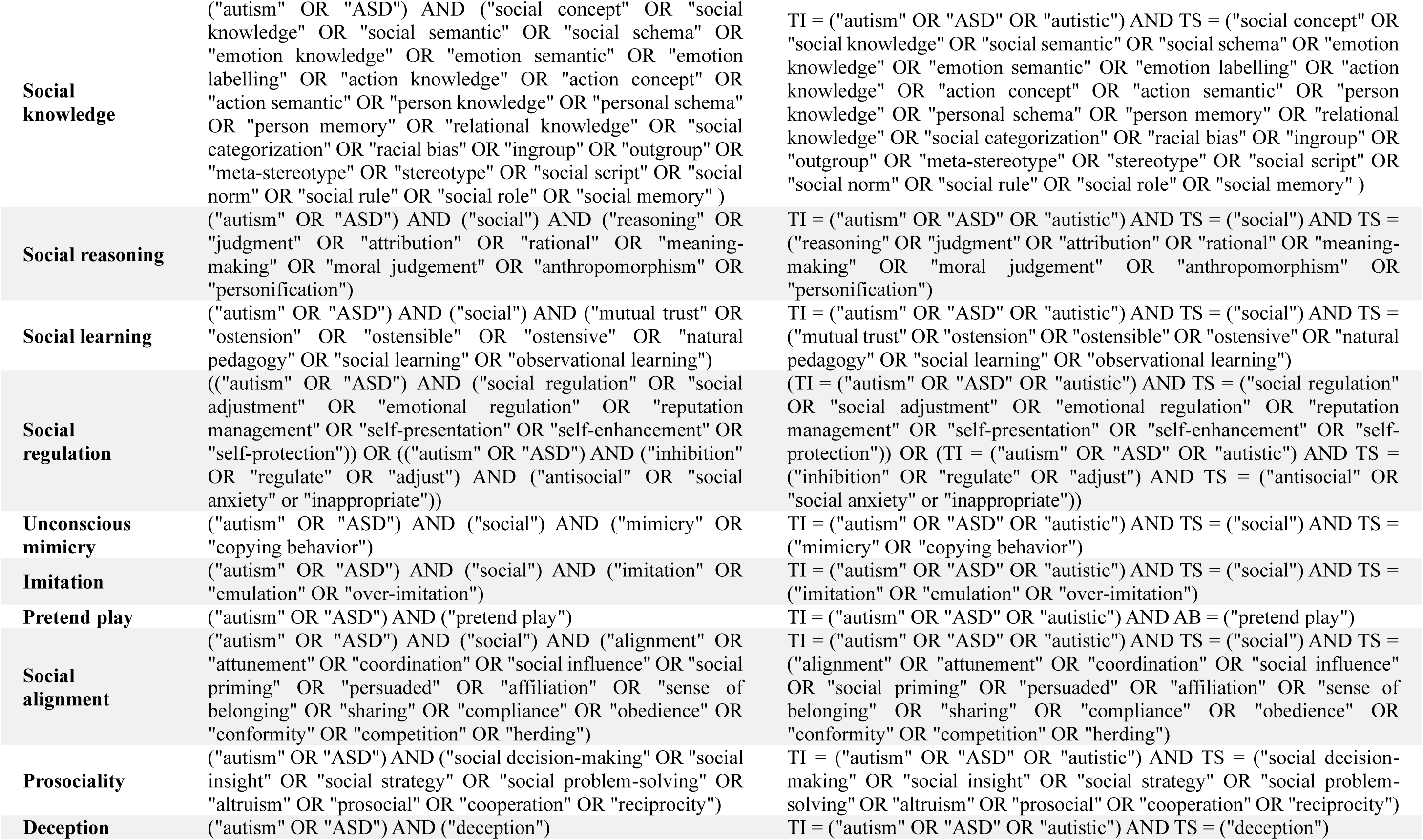
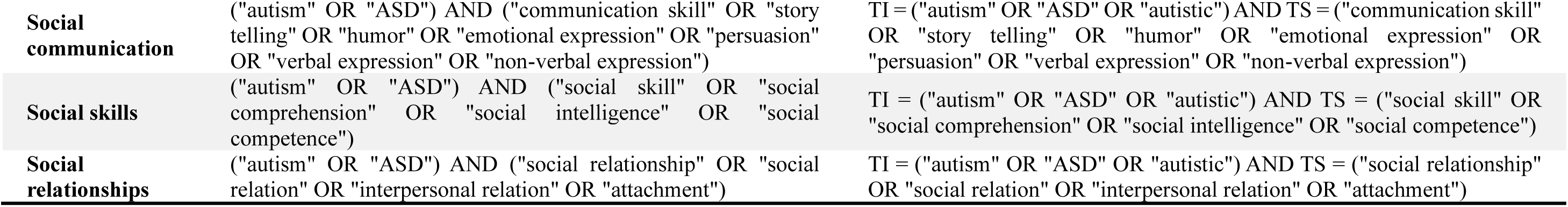
A list of searching terms in PubMed and WoS.

**Supplementary Table 2.**
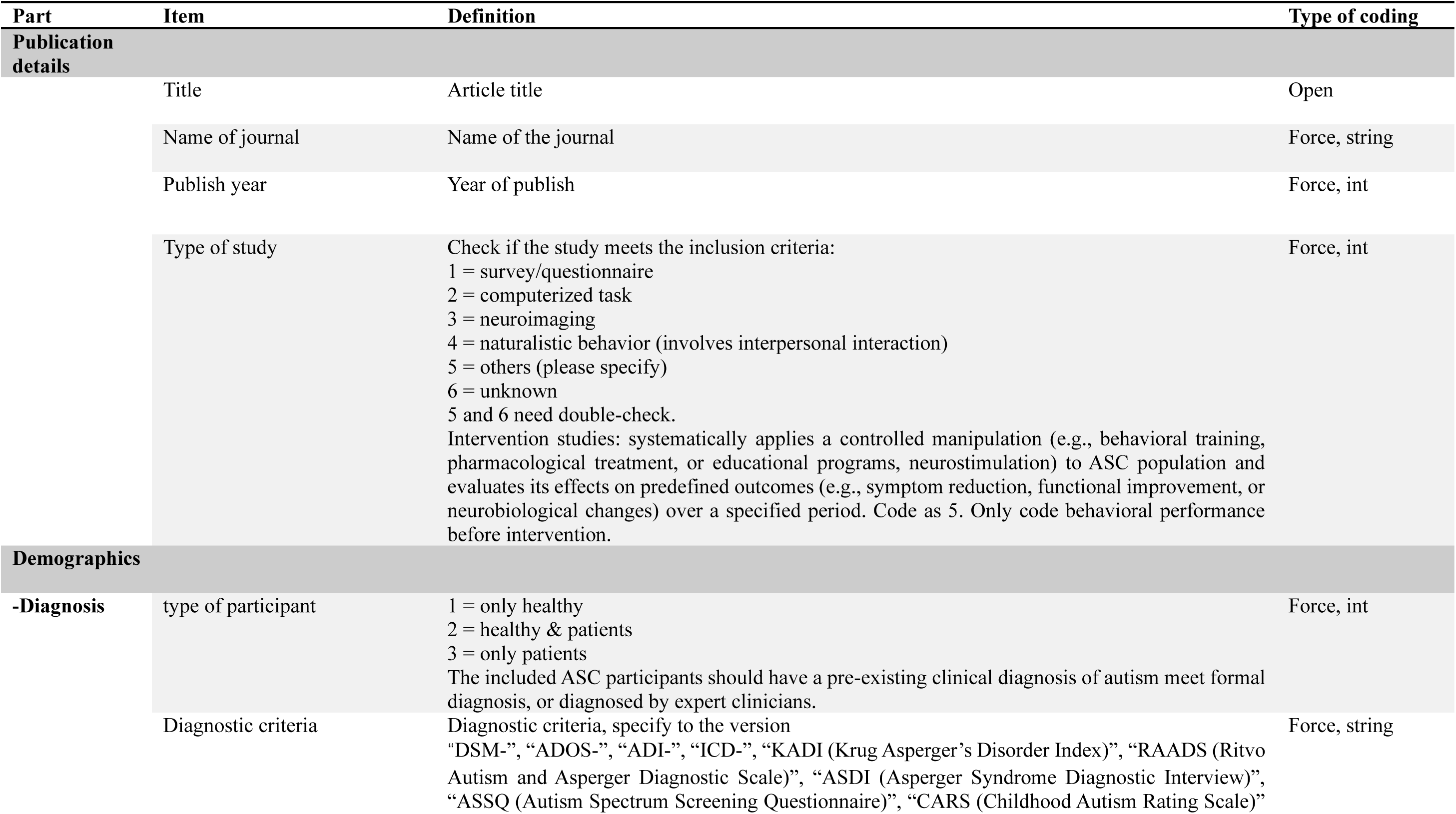
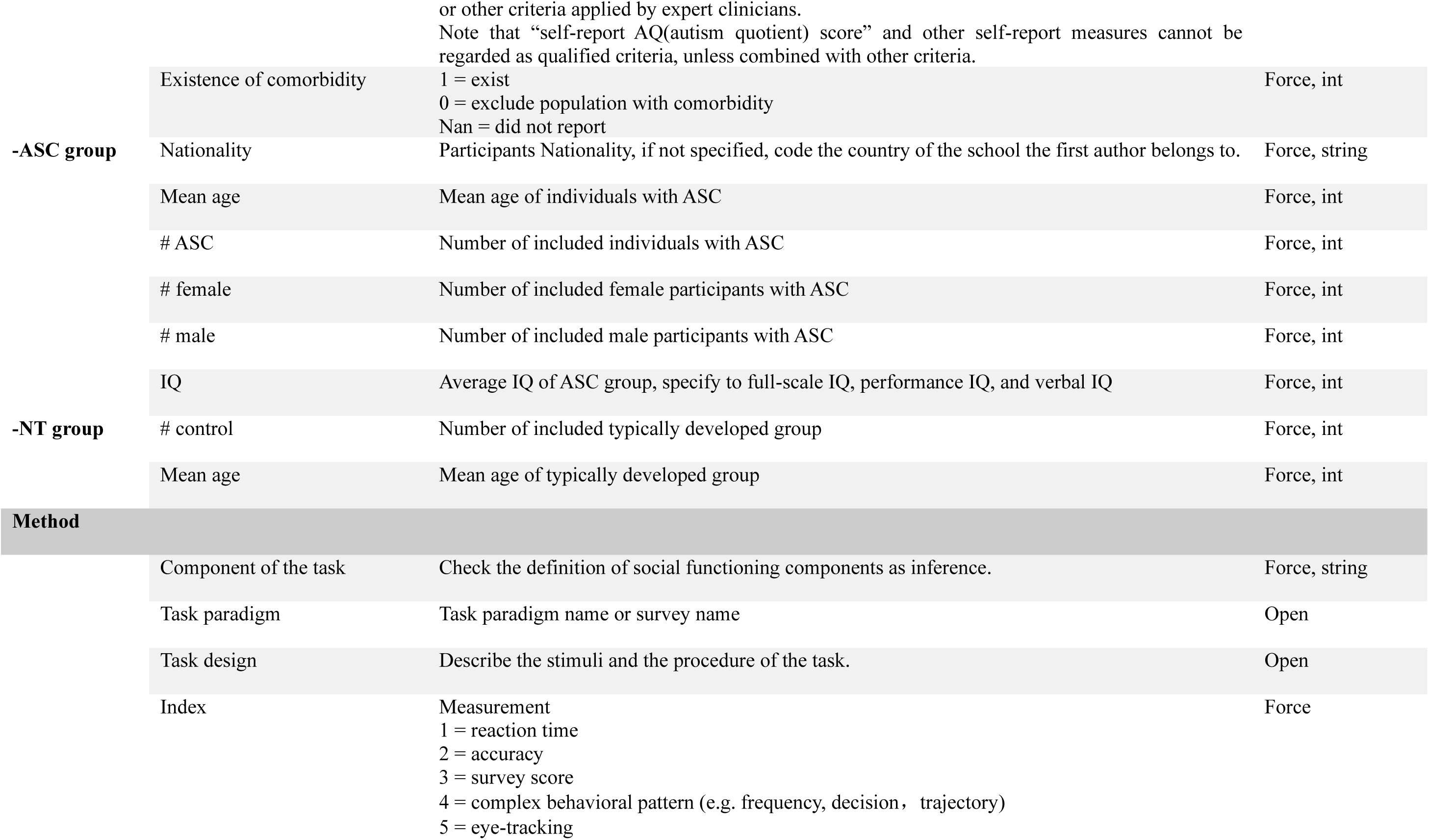
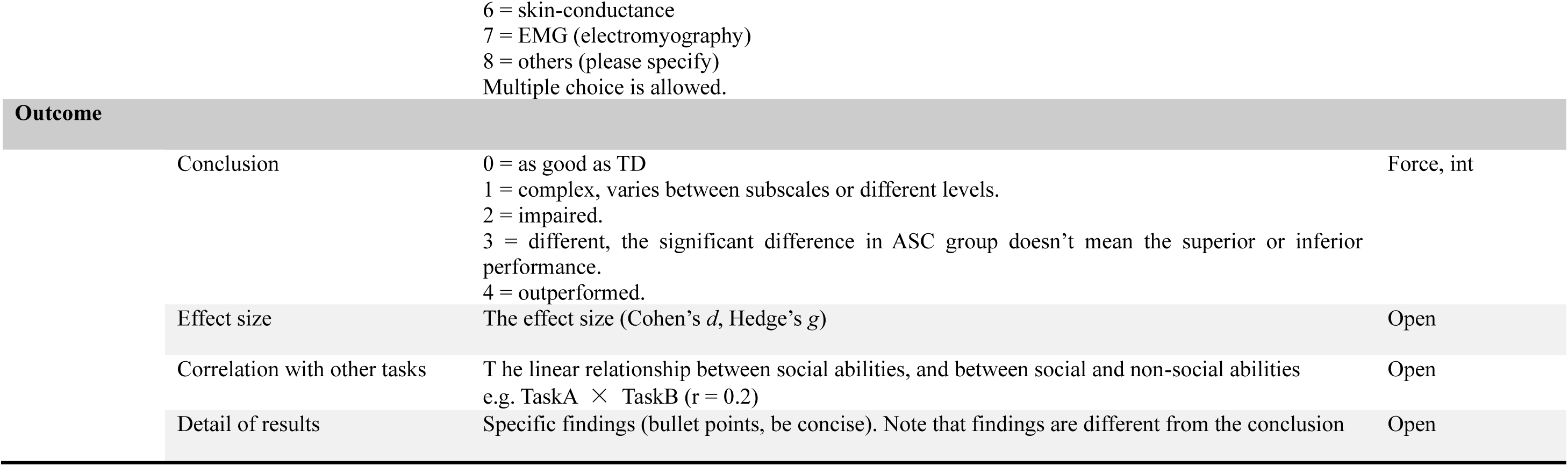
Codebook. There are four parts (publication details, demographics, method and outcome) and two major types of coding (force-coding and open-coding). The inter-coder consistency of force-choice items in 10% articles were double-checked by a third coder, and the consistency rate was 91%.

**Supplementary Table 3.** Ecological validity scoring manual.

|  | Stimuli | Manipulation of stimuli | Feature of stimuli | Data collection | Response |  |
| --- | --- | --- | --- | --- | --- | --- |
| Task | Visual, word, verbal, gesture, 1 point each | Strict-controlled = 0<br>with noises = 1 | Static = 1<br>both = 2<br>dynamic = 3 | non-interactive = 1<br>interactive = 2 | passive perception = 1<br>binary-choice = 2<br>multiple-choice = 3<br>physical movement = 4<br>verbal expression = 5 | Sum |
| Face memory | 1 | 0 | 0 | 1 | 2 | 4 |
| Face discrimination | 1 | 0 | 0 | 1 | 2 | 4 |
| Social reward | 1 | 0 | 0 | 1 | 2 | 4 |
| Social attention: free viewing | 1 | 1 | 0 | 1 | 1 | 4 |
| Facial trait judgement | 1 | 0 | 0 | 1 | 3 | 5 |
| RMET | 1 | 0 | 0 | 1 | 3 | 5 |
| Emotion matching | 1 | 0 | 0 | 2 | 2 | 5 |
| Social seeking | 1 | 0 | 0 | 2 | 2 | 5 |
| Emotion judgment | 1 | 0 | 0 | 2 | 2 | 5 |
| Rating for pain | 1 | 0 | 0 | 1 | 3 | 5 |
| Non-affective plot-light display | 1 | 0 | 0 | 3 | 2 | 6 |
| Emotion labelling | 1 | 0 | 0 | 2 | 3 | 6 |
| Visual perspective-taking | 2 | 0 | 1 | 1 | 2 | 6 |
| Social animation | 1 | 0 | 0 | 3 | 3 | 7 |
| False belief | 2 | 0 | 1 | 2 | 2 | 7 |
| Affective plot-light display | 1 | 0 | 0 | 3 | 3 | 7 |
| Facial mimicry | 1 | 0 | 0 | 3 | 4 | 8 |
| RJA | 2 | 1 | 1 | 3 | 1 | 8 |
| Strange story | 2 | 0 | 1 | 1 | 5 | 9 |
| **Face expression imitation** | 1 | 0 | 1 | 3 | 4 | 9 |
| **Hand/body gesture imitation** | 1 | 0 | 1 | 3 | 4 | 9 |
| **Object imitation** | 1 | 0 | 1 | 3 | 4 | 9 |
| **NEPSY-ToM** | 2 | 0 | 1 | 1 | 5 | 9 |
| **Faux pas** | 2 | 0 | 1 | 1 | 5 | 9 |
| **TASIT** | 2 | 0 | 1 | 1 | 5 | 9 |
| **Initiate joint attention** | 3 | 1 | 1 | 3 | 3 | 11 |
| **Interactive RJA** | 3 | 1 | 1 | 3 | 3 | 11 |
| **Empathy accuracy** | 2 | 1 | 0 | 3 | 5 | 11 |
| **MASC** | 2 | 1 | 1 | 3 | 5 | 12 |

**Supplementary Table 4.**
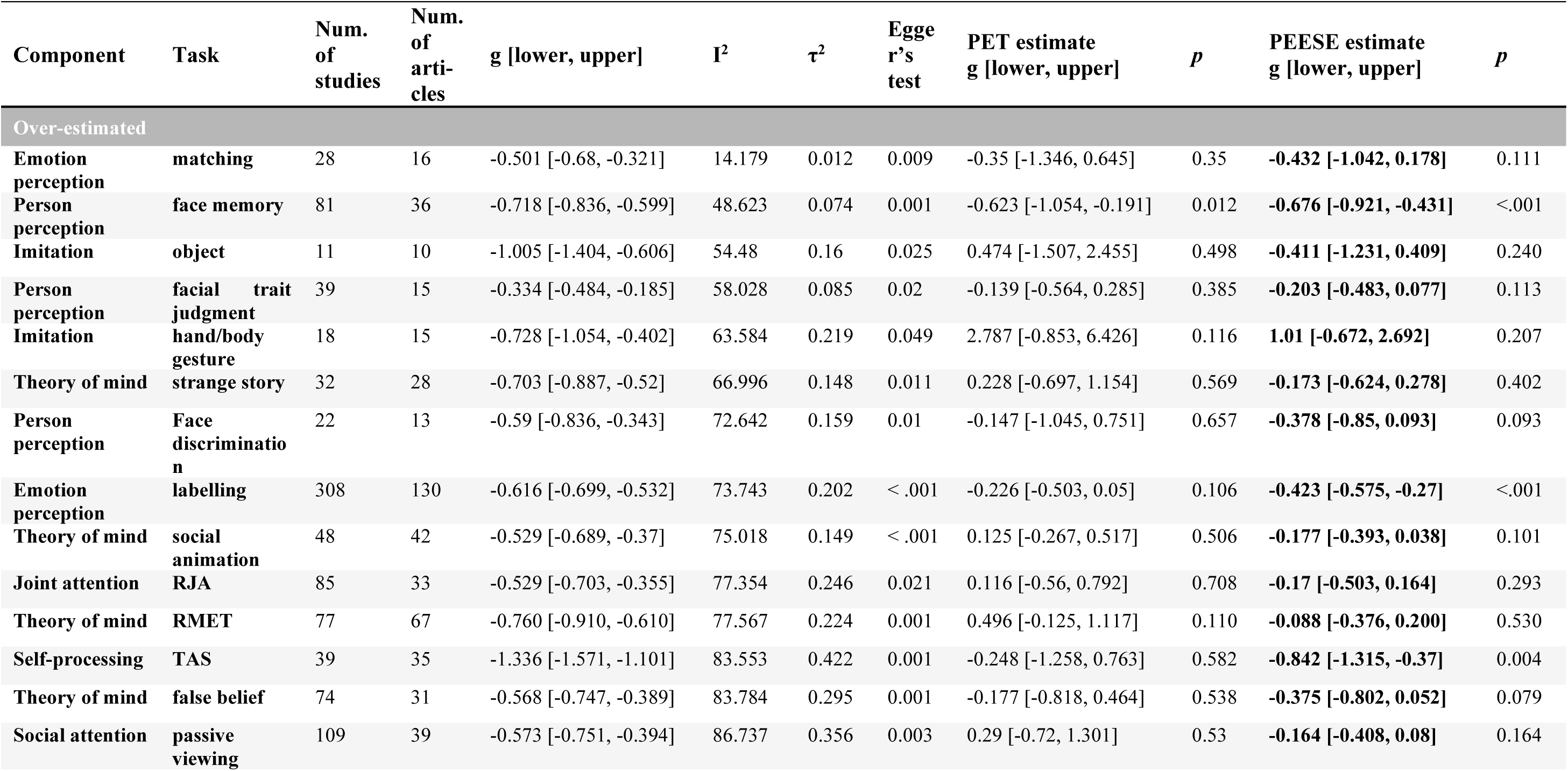
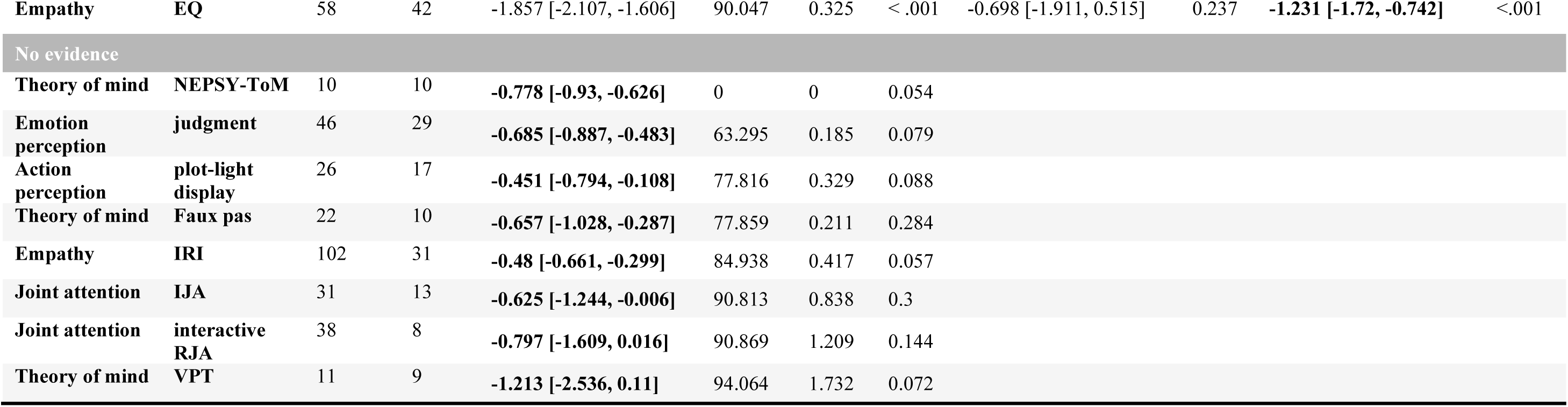
Publication bias analysis for different tasks. Including the number of studies and articles, heterogeneity (I^2^ and τ^2^), *p*-value of asymmetry test (Egger’s test), PET test and adjusted effect sizes, PEESE test and adjusted effect sizes with 95% confidence intervals. “over-estimated” indicated the existence of publication bias, “no evidence” indicated or the egger’s test was not significant. PET test and PEESE estimate was applied for over-estimated tasks. for tasks with substantial heterogeneity (I² > 75%), this adjusted estimate was interpreted as a sensitivity analysis, as asymmetry may stem from true between-study variance rather than publication bias.

| Component | Task | Num. of studies | Num. of articles | g [lower, upper] | $I^2$ | $\tau^2$ | Egger's test | PET estimate g [lower, upper] | $p$ | PEESE estimate g [lower, upper] | $p$ |
| --- | --- | --- | --- | --- | --- | --- | --- | --- | --- | --- | --- |
| Over-estimated |  |  |  |  |  |  |  |  |  |  |  |
| Emotion perception | matching | 28 | 16 | -0.501 [-0.68, -0.321] | 14.179 | 0.012 | 0.009 | -0.35 [-1.346, 0.645] | 0.35 | **-0.432 [-1.042, 0.178]** | 0.111 |
| Person perception | face memory | 81 | 36 | -0.718 [-0.836, -0.599] | 48.623 | 0.074 | 0.001 | -0.623 [-1.054, -0.191] | 0.012 | **-0.676 [-0.921, -0.431]** | <.001 |
| Imitation | object | 11 | 10 | -1.005 [-1.404, -0.606] | 54.48 | 0.16 | 0.025 | 0.474 [-1.507, 2.455] | 0.498 | **-0.411 [-1.231, 0.409]** | 0.240 |
| Person perception | facial trait judgment | 39 | 15 | -0.334 [-0.484, -0.185] | 58.028 | 0.085 | 0.02 | -0.139 [-0.564, 0.285] | 0.385 | **-0.203 [-0.483, 0.077]** | 0.113 |
| Imitation | hand/body gesture | 18 | 15 | -0.728 [-1.054, -0.402] | 63.584 | 0.219 | 0.049 | 2.787 [-0.853, 6.426] | 0.116 | **1.01 [-0.672, 2.692]** | 0.207 |
| Theory of mind | strange story | 32 | 28 | -0.703 [-0.887, -0.52] | 66.996 | 0.148 | 0.011 | 0.228 [-0.697, 1.154] | 0.569 | **-0.173 [-0.624, 0.278]** | 0.402 |
| Person perception | Face discrimination | 22 | 13 | -0.59 [-0.836, -0.343] | 72.642 | 0.159 | 0.01 | -0.147 [-1.045, 0.751] | 0.657 | **-0.378 [-0.85, 0.093]** | 0.093 |
| Emotion perception | labelling | 308 | 130 | -0.616 [-0.699, -0.532] | 73.743 | 0.202 | < .001 | -0.226 [-0.503, 0.05] | 0.106 | **-0.423 [-0.575, -0.27]** | <.001 |
| Theory of mind | social animation | 48 | 42 | -0.529 [-0.689, -0.37] | 75.018 | 0.149 | < .001 | 0.125 [-0.267, 0.517] | 0.506 | **-0.177 [-0.393, 0.038]** | 0.101 |
| Joint attention | RJA | 85 | 33 | -0.529 [-0.703, -0.355] | 77.354 | 0.246 | 0.021 | 0.116 [-0.56, 0.792] | 0.708 | **-0.17 [-0.503, 0.164]** | 0.293 |
| Theory of mind | RMET | 77 | 67 | -0.760 [-0.910, -0.610] | 77.567 | 0.224 | 0.001 | 0.496 [-0.125, 1.117] | 0.110 | **-0.088 [-0.376, 0.200]** | 0.530 |
| Self-processing | TAS | 39 | 35 | -1.336 [-1.571, -1.101] | 83.553 | 0.422 | 0.001 | -0.248 [-1.258, 0.763] | 0.582 | **-0.842 [-1.315, -0.37]** | 0.004 |
| Theory of mind | false belief | 74 | 31 | -0.568 [-0.747, -0.389] | 83.784 | 0.295 | 0.001 | -0.177 [-0.818, 0.464] | 0.538 | **-0.375 [-0.802, 0.052]** | 0.079 |
| Social attention | passive viewing | 109 | 39 | -0.573 [-0.751, -0.394] | 86.737 | 0.356 | 0.003 | 0.29 [-0.72, 1.301] | 0.53 | **-0.164 [-0.408, 0.08]** | 0.164 |
| **Empathy** | **EQ** | 58 | 42 | -1.857 [-2.107, -1.606] | 90.047 | 0.325 | < .001 | -0.698 [-1.911, 0.515] | 0.237 | **-1.231 [-1.72, -0.742]** | <.001 |
| **No evidence** |  |  |  |  |  |  |  |  |  |  |  |
| **Theory of mind** | **NEPSY-ToM** | 10 | 10 | **-0.778 [-0.93, -0.626]** | 0 | 0 | 0.054 |  |  |  |  |
| **Emotion perception** | **judgment** | 46 | 29 | **-0.685 [-0.887, -0.483]** | 63.295 | 0.185 | 0.079 |  |  |  |  |
| **Action perception** | **plot-light display** | 26 | 17 | **-0.451 [-0.794, -0.108]** | 77.816 | 0.329 | 0.088 |  |  |  |  |
| **Theory of mind** | **Faux pas** | 22 | 10 | **-0.657 [-1.028, -0.287]** | 77.859 | 0.211 | 0.284 |  |  |  |  |
| **Empathy** | **IRI** | 102 | 31 | **-0.48 [-0.661, -0.299]** | 84.938 | 0.417 | 0.057 |  |  |  |  |
| **Joint attention** | **IJA** | 31 | 13 | **-0.625 [-1.244, -0.006]** | 90.813 | 0.838 | 0.3 |  |  |  |  |
| **Joint attention** | **interactive RJA** | 38 | 8 | **-0.797 [-1.609, 0.016]** | 90.869 | 1.209 | 0.144 |  |  |  |  |
| **Theory of mind** | **VPT** | 11 | 9 | **-1.213 [-2.536, 0.11]** | 94.064 | 1.732 | 0.072 |  |  |  |  |

**Supplementary Table 5.**
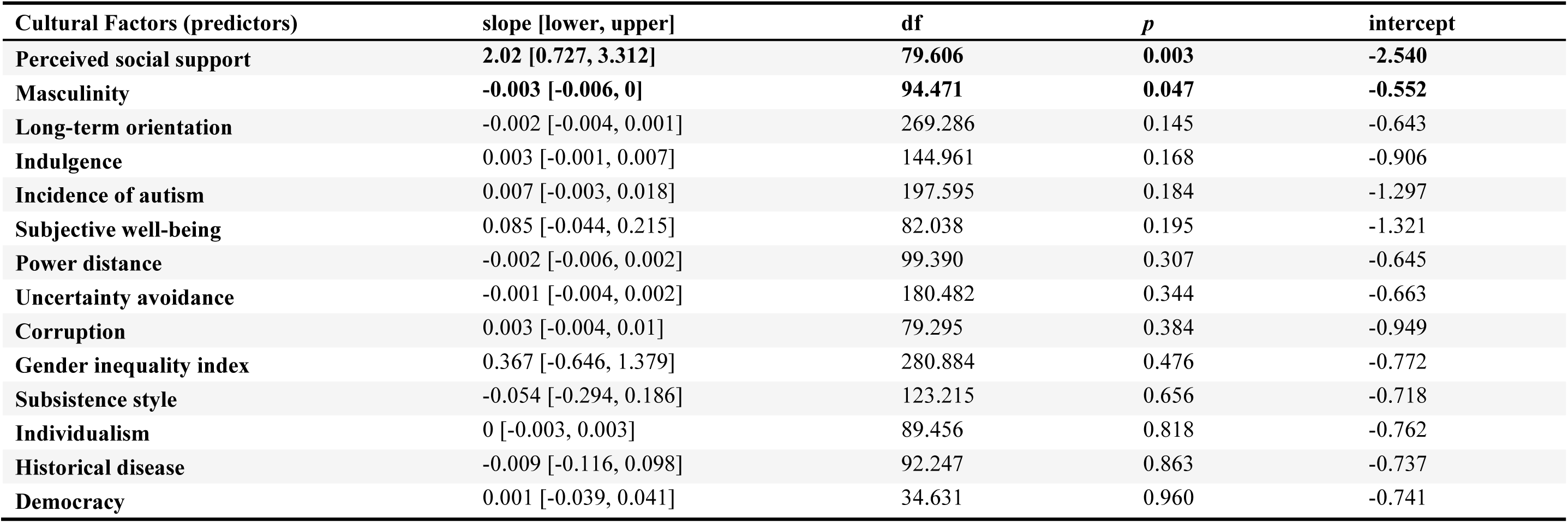
Exploratory meta-regression analysis for sociocultural factors (dependent variable = performance).

**Supplementary Table 6.** Comparison of three competing MASEM models. Estimate = unstandardized parameter estimate; SE = standard error of the estimate; Z = Wald test statistic (Estimate/SE); Pr(>|z|) = two-tailed p-value for the Wald test.

|  |  |  | NT |  |  |  | ASC-NT |  |  |  |
| --- | --- | --- | --- | --- | --- | --- | --- | --- | --- | --- |
|  |  |  | Estimate | SE | Z | Pr(>\|z\|) | Estimate | SE | Z | Pr(>\|z\|) |
| **Serial model** |  |  |  |  |  |  |  |  |  |  |
| **Motor** | → | Emotion | 0.476 | 0.113 | 4.219 | **<.001** | -0.055 | 0.151 | -0.363 | 0.717 |
| **Emotion** | → | Inference | 0.264 | 0.046 | 5.751 | **<.001** | 0.144 | 0.068 | 2.114 | **0.035** |
| **Inference** | → | Complex | 0.226 | 0.041 | 5.545 | **<.001** | 0.160 | 0.066 | 2.428 | **0.015** |
| **Motivation** | → | Motor | 0.175 | 0.068 | 2.578 | **0.010** | 0.193 | 0.141 | 1.369 | 0.171 |
| **Radial model** |  |  |  |  |  |  |  |  |  |  |
| **Motor** | → | Emotion | 0.384 | 0.098 | 3.926 | **<.001** | 0.046 | 0.181 | 0.255 | 0.798 |
| **Emotion** | → | Inference | 0.198 | 0.053 | 3.753 | **<.001** | 0.185 | 0.081 | 2.281 | **0.023** |
| **Inference** | → | Complex | 0.180 | 0.047 | 3.821 | **<.001** | 0.147 | 0.086 | 1.722 | 0.085 |
| **Motivation** | → | Motor | 0.153 | 0.065 | 2.349 | **0.019** | 0.263 | 0.143 | 1.834 | 0.067 |
| **Motivation** | → | Emotion | 0.203 | 0.111 | 1.830 | 0.067 | -0.339 | 0.220 | -1.537 | 0.124 |
| **Motivation** | → | Inference | 0.197 | 0.065 | 3.013 | **0.003** | 0.037 | 0.143 | 0.262 | 0.794 |
| **Motivation** | → | Complex | 0.170 | 0.066 | 2.570 | **0.010** | 0.064 | 0.190 | 0.336 | 0.737 |
| **Network model** |  |  |  |  |  |  |  |  |  |  |
| **Motor** | → | Emotion | 0.350 | 0.086 | 4.056 | **<.001** | 0.071 | 0.172 | 0.409 | 0.682 |
| **Motor** | → | Inference | 0.272 | 0.068 | 3.992 | **<.001** | -0.279 | 0.160 | -1.742 | 0.082 |
| **Motor** | → | Complex | -0.025 | 0.109 | -0.229 | 0.819 | 0.532 | 0.250 | 2.131 | **0.033** |
| **Emotion** | → | Inference | 0.076 | 0.069 | 1.089 | 0.276 | 0.310 | 0.121 | 2.569 | **0.010** |
| **Emotion** | → | Complex | 0.082 | 0.098 | 0.837 | 0.403 | -0.042 | 0.228 | -0.182 | 0.855 |
| **Inference** | → | Complex | 0.173 | 0.058 | 2.995 | **0.003** | 0.070 | 0.134 | 0.525 | 0.600 |
| **Motivation** | → | Motor | 0.132 | 0.064 | 2.063 | **0.039** | 0.234 | 0.142 | 1.656 | 0.098 |
| **Motivation** | → | Emotion | 0.223 | 0.107 | 2.080 | **0.038** | -0.378 | 0.220 | -1.721 | 0.085 |
| **Motivation** | → | Inference | 0.191 | 0.065 | 2.939 | **0.003** | 0.090 | 0.180 | 0.501 | 0.616 |
| Motivation | → | Complex | 0.144 | 0.072 | 1.996 | **0.046** | -0.168 | 0.243 | -0.691 | 0.489 |

